# 3D matrix adhesion composition facilitates nuclear force coupling to drive invasive cell migration

**DOI:** 10.1101/2021.05.17.443835

**Authors:** Daniel Newman, Lorna Young, Thomas Waring, Louise Brown, Katarzyna Wolanska, Ewan Macdonald, Arthur Charles Orszag, Patrick Caswell, Tetsushi Sakuma, Takashi Yamamoto, Laura Machesky, Mark Morgan, Tobias Zech

## Abstract

Cell invasion and metastasis is a multi-step process, initialised through the acquisition of a migratory phenotype and the ability to move through differing and complex 3D extracellular environments. In this study we set out to identify the parameters required for invasive cell migration in 3D environments. Cells interact with the extracellular matrix via transmembrane-spanning integrin adhesion complexes, which are well characterised in cells plated on 2D surfaces, yet much less is known about them in cells embedded in 3D matrices. We establish a technique to determine the composition of cell matrix adhesion complexes of invasive breast cancer cells in 3D matrices and on 2D surfaces and we identify an interaction complex enriched in 3D adhesive sites required for 3D invasive migration. Depletion of β-PIX-Myosin18A (Myo18A) abolishes cancer cell invasion, without negatively affecting matrix degradation, Rho GTPase signalling, or protrusion formation in collagen matrices. Instead, in a mechanism only seen in cells moving through 3D matrix, β-PIX and Myo18A drive the polarised recruitment of non-muscle Myosin 2A (NM2A) to the tips of protrusions. This recruitment of NM2A is required for the creation of an NM2A-NM2B isoform gradient, which ranges from the protrusion to the nucleus. We observe a requirement for active force transmission to the nucleus during invasive migration that is needed to pull the nucleus forward. We postulate that the establishment of the NM2A-NM2B actomyosin gradient facilitates the coupling of cell-matrix interactions at the protrusive cell front with nuclear movement, enabling effective invasive migration and front-rear cell polarity.

## Introduction

Cell migration is fundamental for normal physiology and is dysregulated in many diseases, including cancer. Efficient migration depends critically on how cells sense and interact with their surrounding microenvironment and regulate cytoskeleton dynamics (Caswell and Zech, 2018; van Helvert et al., 2018). Migrating cells interpret and influence the physical and biochemical properties of the extracellular matrix (ECM) as well as interacting with neighbouring/proximal cells. The mechanical and compositional properties of ECM are detected primarily by membrane spanning integrin-associated adhesion complexes (IACs). IACs act as bidirectional signalling hubs, forming a mechanical link between the cytoskeleton and ECM. A large number of proteins dynamically associate with IACs, changing over the lifecycle of adhesion complexes and in response to their microenvironment. IACs formed on planar 2-dimensional (2D) substrates have been extensively studied using numerous techniques including super-resolution and live-cell microscopy, as well as biochemical isolation techniques (Ajeian et al., 2016; Chastney et al., 2020; Humphries et al., 2009; Jones et al., 2015; Kuo et al., 2011; Schiller et al., 2011). These studies have provided great insight into the composition and regulation of IACs in 2D, however biochemical isolation techniques are not transferrable to cells in 3D culture.

There are few situations *in vivo* where cells encounter a 2D environment. Instead, most cells are surrounded and functionally integrated with 3-dimensional (3D) ECM. A 3D microenvironment presents a number of challenges to migrating cells that are not present on 2D substrates, including physical and topological cues and barriers. Compared with IACs on 2D substrates, little is known about the composition, regulation and role of IACs in cells embedded in 3D microenvironments but initial analysis of IACs in 3D matrix and *in vivo* suggested that 3D IACs exhibit differences in composition, and are smaller in size in comparison with their 2D counterparts (Cukierman et al., 2001; Fraley et al., 2010; Harunaga and Yamada, 2011). Importantly, invasive cancer cells have hybrid adhesion structures in 3D; containing both the adhesion marker paxillin and the actin polymerisation factor N-WASP, often associated with invadopodial structures on 2D substrates (Yu et al., 2012). To date, technological limitations have prevented comprehensive analysis of IAC signalling networks in 3D matrix.

IACs link the extracellular matrix to the cytoskeleton. Adaptor proteins like talin and vinculin can bind both adhesion receptors and the actin cytoskeleton (Goult et al., 2018; Seetharaman and Etienne-Manneville, 2020). IACs are involved in organising actin into contractile actin networks containing non-muscle Myosin 2 (NM2) bundles (Tojkander et al., 2012; Vignaud et al., 2021). In addition, IACs have several connections to actin polymerisation factors through recruitment of GEFs (Müller et al., 2020), WASP family members, and formins (Alieva et al., 2019; Ciobanasu et al., 2012; Livne and Geiger, 2016). The actin arrangement associated with newly seeded IACs is not well characterised and has been described as an actin “cloud” potentially polymerised by the transient recruitment of Arp2/3 complex through FAK and vinculin (DeMali et al., 2002; Serrels et al., 2007). Mature adhesion complexes tend to be functionally connected to actomyosin stress fibres, whereas early/nascent adhesion have less associated NM2 (Tojkander et al., 2012). The mechanism of this switch to NM2 association is currently unknown. Many cells express several NM2 isoforms with unique motor parameters and distinct functions (Barua et al., 2014; Kovács et al., 2007; Kovács et al., 2003). NM2 isoforms are not uniformly distributed throughout the cell, but exhibit a polarised distribution (Vicente-Manzanares et al., 2011). Shutova and colleagues recently proposed that the prevalent recruitment from non-muscle Myosin 2A () to cell periphery is driven by its higher expression level whereas the concentration of non-muscle Myosin 2B (NM2B) towards the cell interior is due to its stronger binding to polymerised actin (Shutova et al., 2017). However, the function of the NM2 isoform gradient or its impact on invasive cell migration is unclear.

A major barrier to successful 3D migration is the ability of cells to move their nuclei through geometrically restrictive, densely cross-linked ECM (Davidson and Lammerding, 2014; Harada et al., 2014; Thomas et al., 2015; Wolf et al., 2013). The outer nuclear membrane is coupled to the cytoskeletal network through the Linker of Nucleoskeleton and Cytoskeleton (LINC) (Hieda, 2019). The LINC complex contains the Nesprin family of proteins, which connect to cytoskeletal components, and SUN domain proteins, which connect to the inner nuclear lamina. The complex therefore acts as a mechanical conduit to bridge the outer-nuclear-to the inner-nuclear-cytoskeleton. Consequently, a functional LINC complex is critical for cell migration (Lee et al., 2007; Thomas et al., 2015), nuclei movement, positioning and shape (Arsenovic et al., 2016; Horn et al., 2013; Luxton et al., 2010), centrosome function and positioning (Chang et al., 2015) and is involved in force sensitive transcriptional pathways (Bouzid et al., 2019).

In spite of the wealth of knowledge about cell migration, it is currently unclear how cells are able to couple nuclear movement with migration, and importantly, what role IACs may play in this process, particularly during 3D migration. To interrogate this, we developed an approach to directly compare IAC composition and signalling networks on 2D and in 3D extracellular matrix using a BioID2-based proximity-labelling coupled with proteomics (Kim et al., 2016). We identify proteins in breast cancer cells that have not previously been associated with adhesions, a large number of proteins that are less abundant in 3D adhesions and, critically, a defined group that exhibited enhanced enrichment in 3D adhesions, including βPix and Myosin18A (Myo18A). Downstream analysis revealed that βPix and Myo18A are essential for efficient invasion without affecting the ability of the cell to form protrusions or degrade matrix. By contrast, βPix and Myo18A control adhesion maturation and the generation of the non-muscle myosin-2 (NM2A and NM2B) isoform-gradient via recruitment of NM2A to the leading edge of protrusions. Failure to establish the membrane-nuclear NM2A/B gradient, due to perturbation at adhesions or the LINC complex, leads to loss of nuclear force coupling, front-rear polarity and 3D invasive migration.

## Results

### Identification of cell matrix adhesion networks in 3D environments

Previous proteomic studies have provided comprehensive systematic analysis of IACs, predominantly from fibroblast cells, plated on 2D matrix substrates (Horton et al., 2016). These analyses led to the definition of the “consensus adhesome”, a core set of proteins recruited to IACs, which were sub-divided into four distinct signalling modules (Horton et al., 2016). However, these proteomic strategies have not been used to analyse the composition of adhesions in invasive cancer cells embedded in 3D matrices. To investigate the composition of IACs in invasive triple negative breast cancer cells (MDA-MB-231), we developed an approach to compare adhesion protein complexes from 2D versus 3D extracellular matrices. We utilised a promiscuous biotin ligase-based proximity-dependent labelling approach, BioID2, which can label proteins within approximately 10-20 nm range of the bait protein (Kim et al., 2016). We generated two different BioID2 bait constructs fused to the consensus adhesome proteins paxillin (BioID2-Paxillin) and talin (Talin--BioID2) (Figure 1A)(Horton et al., 2016). Paxillin and talin were chosen as both are known to act as scaffolding proteins and are associated with nascent as well as mature adhesions (Brown and Turner, 2004; Klapholz and Brown, 2017). Importantly, the respective N- and C-termini of the two proteins have been reported to be localised at different axial strata within 2D adhesion complexes. Paxillin localises close to the plasma membrane, whereas the C-terminus of talin is positioned proximal to actin stress fibres (Kanchanawong et al., 2010).

**Figure 1.**
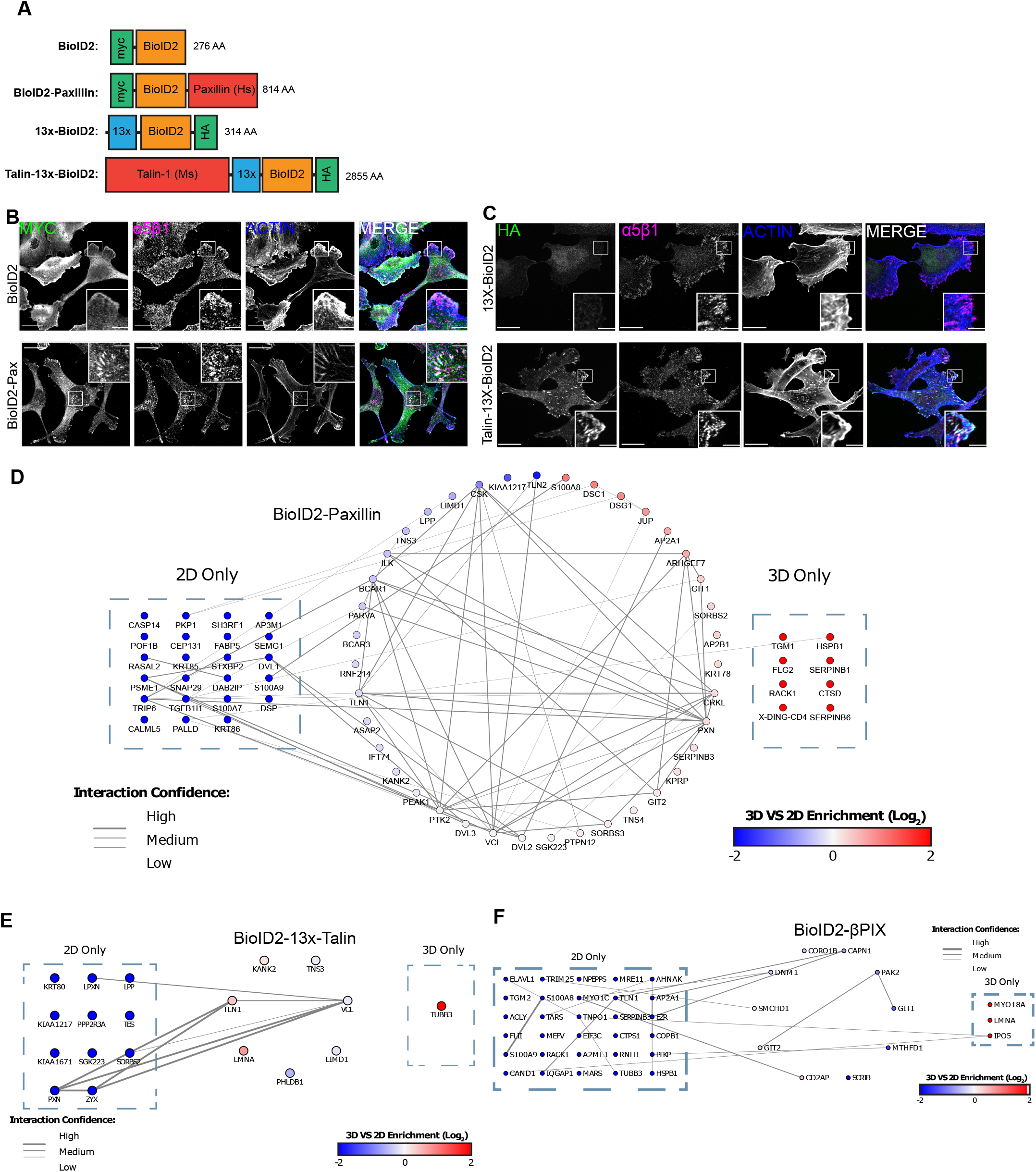
Comparison of IAC-associated proteins in MDA-MB-231 cells cultured in a 2D vs 3D microenvironment. (A) Schematic of BioID2 adhesion constructs used in this study. (B) Immunostaining of MDA-MB-231 cell lines stably expressing BioID2 and BioID2-Paxillin on 2D substrates. Scale bar in expanded view = 20 µm, in magnified view = 5 µm. (C) Immunostaining of MDA-MB-231 cells stably expressing 13x-BioID2 or Talin-13x-BioID2. Scale bar in expanded view = 20 µm, in magnified view = 5 µm. (D & E) Protein-protein interaction network showing all proteins identified as being enriched in BioID2-Paxillin (D) or BioID2-13x-Talin (E) compared with corresponding BioID2 control (Fisher-Exact, P < 0.05 Benjamini-Hochberg corrected) (n=3). Interaction network obtained from STRING 10.5 (Experimentally identified interactions only, confidence level > 0.4). Protein enrichment is shown based on total spectral counts normalised to BioID2 identification in each condition. (F) Protein-protein interaction network showing all proteins identified as being enriched in BioID2-βPIX. Proteins were considered to be positively enriched if 3 or more spectra were identified and they were enriched 2 fold vs a BioID only expressing conditional control. Proteins identified in both 2D and 3D conditions are coloured to represent 3D vs 2D total spectral counts normalised to total βPIX spectra for that given condition. Interaction network obtained from STRING (10.5) based on interaction database and experimental data. Edge weight represents interaction confidence score. (High > 0.95, Medium 0.95-0.9, Low 0.9-0.4) (n = 1).

MDA-MB-231 cells stably expressing BioID2, BioID2-Paxillin and Talin-13x-BioID2 were generated and verified (Figure 1B, C, Supplementary Figure 1A, B). Cells were cultured either on 2D conditions: plastic dishes coated with fibronectin and collagen, or in 3D conditions: collagen hydrogels supplemented with fibronectin.

Following biotin treatment proteins were extracted, the biotinylated proteins enriched, analysed by LC-MS/MS and subjected to statistical and network analysis. Within the BioID2-Paxillin dataset, a total of 74 proteins (2D: 63, 3D: 51) were identified as significantly enriched across all three repeats compared to the BioID2 only control (Figure 1D). 23 proteins were uniquely identified in 2D and 11 were only present in 3D, leaving 40 proteins which were identified in both conditions. By contrast, 20 proteins were identified in the Talin1-13x-BioID2 dataset (2D: 18, 3D: 9, Figure 1E). 11 were identified in 2D and 2 proteins were present only in 3D, while 7 proteins were identified in both conditions.

The biophysical properties of the cellular microenvironment are known to regulate gene expression. To examine whether the observed changes in protein detection between 2D and 3D culturing conditions were a result of changes in gene expression, RNA-Seq analysis was conducted on cells cultured using the same substrate conditions as those used in the proteomic datasets. Overall, no significant correlation was found between 3D vs. 2D gene expression in RNA-Seq analysis and 3D vs. 2D enrichment of the identified hits in the BioID2-Paxillin or Talin1-13x-BioID2 datasets (Supplementary Figure 1C-E). Seven proteins identified in the BioID2 datasets were found to have significant expression changes in the RNA-seq dataset, with only 4 proteins (RASAL2, HSPB1, SERPINB1, JUP) showing substantial changes in the same direction in both studies. This suggests that for the most part, within the BioID2 datasets, the changes in enrichment observed were largely independent of gene expression changes.

To estimate total coverage of adhesion components, we compared the positively identified proteins within our datasets against the consensus adhesome, which was based on IAC enrichment approaches on 2D fibronectin substrates (Horton et al., 2016). The BioID2-Paxillin dataset generated here achieved identification of 80% of the Paxillin-FAK axis, as well as 80% of the Talin-Vinculin axis (Supplementary Figure 1F). Two additional consensus adhesome proteins were identified in the Talin1-13x-BioID2 dataset (Testin & Zxyin) that were not identified in the BioID2-Paxillin dataset. Both of these proteins form part of the Actinin-Zxyin module.

### βPix and Myosin-18A (Myo18A) enrich at 3D matrix adhesion sites and are required for invasive migration

Having characterised proteins associated with paxillin or talin on 2D or in 3D substrates, we sought to identify candidate regulatory proteins recruited to adhesions in 3D matrix. To achieve this, we focused on the BioID2-Paxillin dataset, as paxillin is located within the adhesion signalling layer of IACs (Kanchanawong et al., 2010), and interrogated the datasets to identify sub-networks of known interactors, each exhibiting a similar enrichment profile towards BioID2-paxillin in 3D ECM. ARHGEF7 (also known as βPix), GIT1 and GIT2 were found to be enriched within 3D matrices in the BioID2-Paxillin dataset (Figure 1D). βPix and the GIT proteins are known to form a complex and are established IAC components in cells cultured on 2D substrates (Kuo et al., 2011; Nayal et al., 2006; Zhao et al., 2000). βPix is also upregulated in breast cancer (Ahn et al., 2003), thus we chose to investigate the role of βPix in 3D embedded cells further. To do so, a BioID-βPix screen in MDA-MB-231 cells was conducted comparing interactors of βPix in 2D versus 3D environments (Figure 1F). This screen identified two proteins that βPix is known to bind: SCRIB and Myo18A (Hsu et al., 2010; Hsu et al., 2014). Of these, only Myo18A was enriched in the 3D matrices compared to the 2D substrate (Figure 1F). These data suggested a role for the βPix-Myo18A complex specifically in 3D microenvironments. Thus, we sought to investigate the potential role of βPix-Myo18A in 3D invasive cell migration.

Initially, βPix and Myo18A localisation within 2D-plated cells was examined. In these conditions, βPix clearly localised at paxillin-positive adhesion sites, where βPix shows accumulation at the membrane-proximal region of adhesion complexes (Figure 2A, B, Supplementary Fig 2A). We were unable to identify an antibody that faithfully stained endogenous Myo18A (data not shown), therefore we examined its localisation through GFP-Myo18A expression with mCherry-Vinculin (Supplementary Figure 2B). We observed accumulation of GFP-Myo18A in adhesions, yet with a more dispersed pattern compared with βPix localisation. Together these data suggest that both βPix and Myo18A can localise at, or in close proximity to, adhesions in MDA-MB-231 cells plated on 2D matrices, as previously reported for βPix in other cell types (Kuo et al., 2011; Oh et al., 1997).

**Figure 2.**
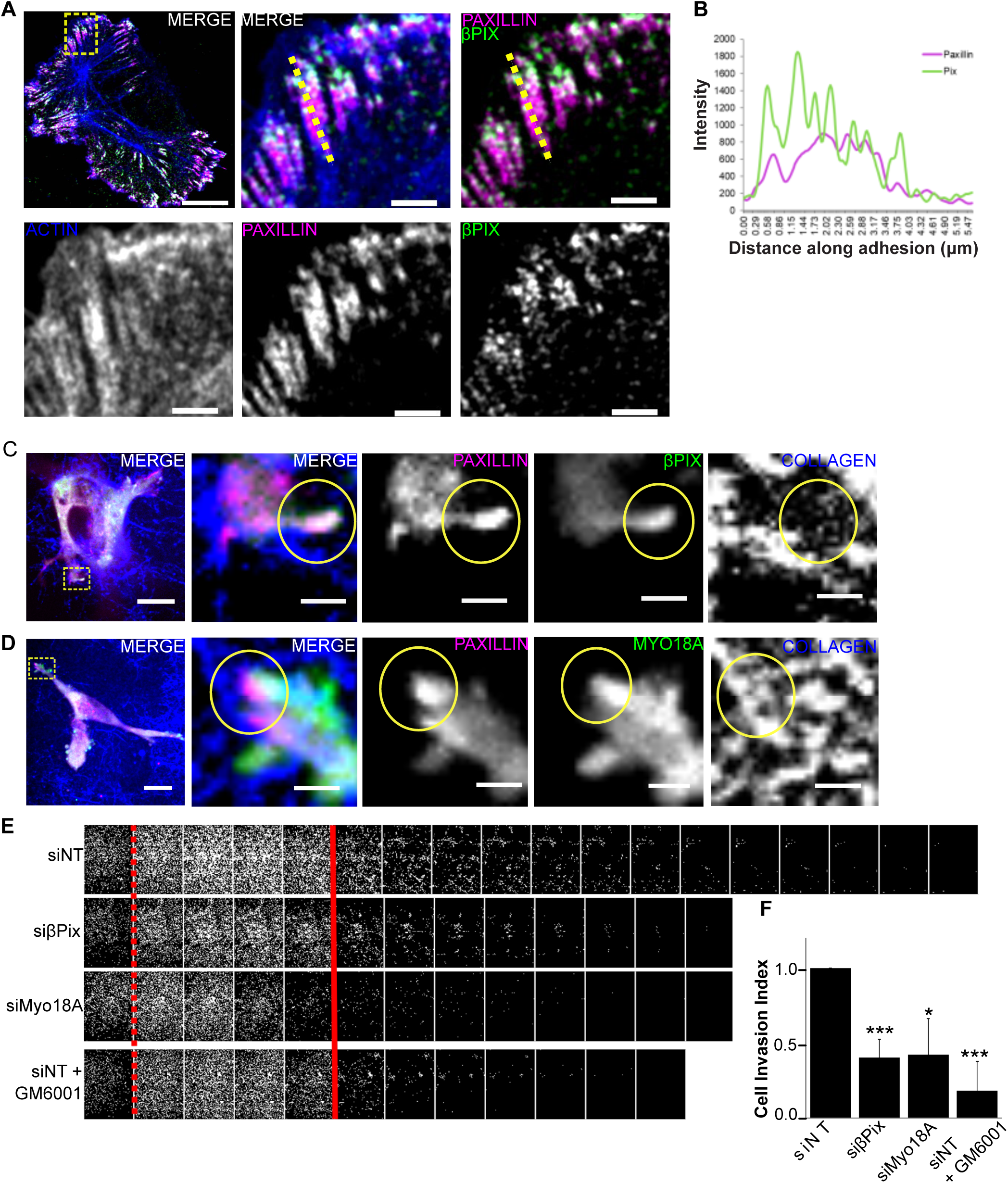
βPix and Myo18A localise to 3D adhesive sites and are required for cell invasion. (A) Representative image of MDA-MB-231 cells fixed and stained with phalloidin (blue), Paxillin (magenta), and βPix (green). Boxed region in expanded view indicates region of magnified view. Images are single Z slice projections. Scale bar in expanded view = 10 µm, in magnified view = 2 µm. (B) Line scan profile of Paxillin and βPix across focal adhesion shown in A with dashed line. Magenta line indicates Paxillin intensity, green line indicates βPix intensity. (C) Representative image of MDA-MB-231 cells transfected with mApple-Paxillin (magenta) and GFP-βPix (green) embedded in collagen (supplemented with ∼1-2% fluorescent collagen (blue) & fibronectin gel). Boxed region in expanded view indicates region of magnified view, circles in magnified view indicate Paxillin/βPix enrichment. Scale bar in expanded view = 10 µm, in magnified view = 2 µm. (D) Representative image of MDA-MB-231 cells transfected with mApple-Paxillin (magenta) and GFP-Myo18A (green) embedded in collagen (supplemented with ∼1-2% fluorescent collagen (blue) & fibronectin gel). Boxed region in expanded view indicates region of magnified view, circles in magnified view indicate Paxillin/Myo18A enrichment. Scale bar in expanded view = 10 µm, in magnified view = 2 µm. (E) Inverted transwell invasion assay. MDA-MB-231 cells treated with non-targeting- (NT), βPix-, or Myo18A-siRNA, or GM6001 (MT1-MMP inhibitor). Cells were subjected to inverted invasion assay in matrigel/fibronectin gels for 5 days. Cells were fixed and stained with Hoesct. Images were taken every 10 μm into gel. (F) Data represents cell invasion index: fold change in percentage of cells invading ≥ 40 μm depth (red solid line)/ N = ≥ 4 experiments. Two sample t-test was used to estimate p values: * p < 0.05, *** p < 0.001. Error bars are SD.

Next, βPix and Myo18A localisation was examined in cells embedded in 3D matrices. Both GFP-βPix and GFP-Myo18A showed enrichment at mApple-Paxillin and mCherry-Vinculin positive adhesion structures which were coincident with collagen fibres (Fig 2C, D, Supplementary Figure 2C, D). This established that βPix and Myo18A are both enriched at adhesive sites in 3D embedded cells.

Having identified that βPix and Myo18A associate with adhesions in 3D matrices we examined what role they may play in cancer cell invasion. The ability of MDA-MB-231 cells to invade extracellular matrices was examined using inverted invasion assays. Cells were embedded in matrigel complemented with fibronectin and examined for their ability to invade into matrices. Depletion of either βPix or Myo18A by siRNA (Supplementary Figure 2E) resulted in a significant decrease in invasion similar to the levels with GM6001 treatment (metalloproteinase inhibitor) (Figure 2E-F).

Effective invasive cell migration through restrictive matrices requires the ability to remodel and degrade the surrounding matrix (Caswell and Zech, 2018; Wolf et al., 2013). To measure the ability of MDA-MB-231 cells, depleted of βPix or Myo18A, to degrade matrices we employed gelatin-based invadopodia assays. Surprisingly, loss of βPix or Myo18A caused an increase in the amount of degradation, measured by percentage of cells associated with degradation spots, as well as degraded patches per cell (Supplementary Figure 2H-J). These results suggest there is no defect in the ability of βPix or Myo18A depleted cells to degrade extracellular matrices.

To examine the effect of βPix or Myo18A depletion on cell migration we performed wound healing assays on cells plated on fibronectin/collagen coated plastic. Depletion of either βPix or Myo18A led to a reduction in wound closure over 40 h by 48% and 78%, respectively, compared to control cells (Supplementary Fig 2F-G).

On 2D surfaces effective cell migration requires precise co-ordination of adhesion dynamics, and βPix has previously been reported to negatively regulate adhesion maturation and promote nascent adhesion turnover (Kuo et al., 2011). Therefore we examined the effect of βPix or Myo18A depletion on matrix adhesions in MDA-MB-231 cells plated on 2D surfaces. Depletion of either protein causes a significant increase in the number of small adhesions (≤ 2 μm squared), and an increase in the number of adhesions per cell (Supplementary Figure 3A-D).

Together these results suggest that βPix and Myo18A are required for efficient migration on 2D surfaces and invasion in 3D matrices. Adhesions are still assembled in βPix and Myo18A depleted cells on 2D but differ in size and number from their wildtype counterparts. In 3D, βPix or Myo18A depletion does not negatively affect matrix degradation, yet does inhibit invasive migration, suggesting a direct impact on cell motility.

### βPix and Myo18A regulate nuclear morphology and internal cell polarity

To further investigate the cause of the migration and invasion defects seen with βPix or Myo18A depletion we examined a variety of parameters related to cell migration. MDA-MB-231 cells form characteristic pseudopodia during 3D cell invasion (Yu et al., 2012), a process that is highly dependent upon actin dynamics. Therefore, the ability of βPix- or Myo18A-depleted cells to assemble actin protrusions in 3D matrices was examined. Following βPix- or Myo18A-depletion no defect in actin protrusion assembly was observed (Supplementary Figure 4A, B), showing that the invasion defect is not due to a defect in global actin assembly parameters.

βPix has been reported to exhibit guanine nucleotide exchange factor (GEF) activity for CDC42 and Rac *in vitro* (Manser et al., 1998), yet this activity is not consistently observed in cells (Müller et al., 2020). We therefore chose to examine global activity of the Rho GTPases Rac1 and CDC42 following βPix depletion using two separate approaches: FRET/FLIM imaging of live-cells expressing Rac1 or Cdc42 FRET biosensors (Hodgson et al., 2010; Itoh et al., 2002), and PAK-PBD pull-down assays to measure Rac1 and Cdc42 activity. No change in Rac1 or Cdc42 activity was observed in βPix-depleted cells in either assay (Supplementary Figure 4C-H).

A reduction in actomyosin contractility could also lead to defective migration (Hiroyasu et al., 2017). Evaluation of MLC phosphorylation in βPix depleted cells did not show a reduction of MLC phosphorylation (Supplementary Figure 4I-J). In addition, RNA-sequencing analysis and western blot analysis suggest that expression levels of NM2A/B heavy chain are unaltered following βPix/Myo18A depletion (Supplementary Figure 5A, B). These data, together with the ability to form pseudopodial protrusions, suggest that the observed migration defects in response to βPix depletion are not a consequence of disruption to Rac and CDC42 signalling or global contractility.

Another factor required for efficient migration is correct nuclear morphology and positioning (Arsenovic et al., 2016; Lee et al., 2007; Luxton et al., 2010; Thomas et al., 2015), therefore the nuclei morphology of cells depleted of βPix or Myo18A was examined in cells embedded in 3D ECM. Unlike their wild-type counterparts which often contained elongated nuclei, oriented along in the axis of the pseudopods, βPix/Myo18A-depletion led to a higher degree of nuclear roundness, and the nuclei were not orientated in the direction of the associated pseudopod (Figure 3 A, B).

**Figure 3:**
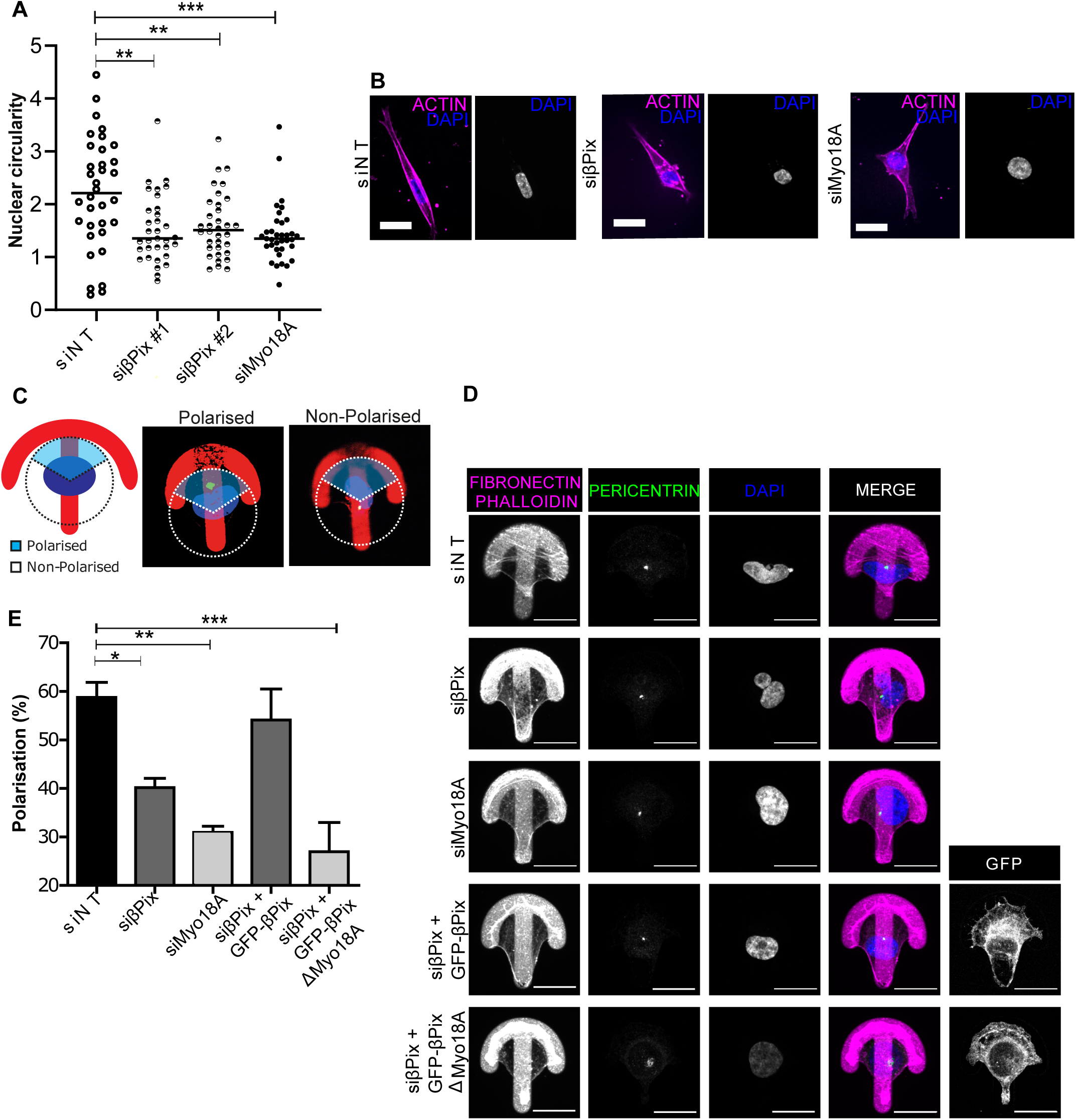
βPix and Myo18A are required for correct nuclear morphology and internal cell polarity. (A) Dot plot shows nuclear index (long/short length) of NT-, βPix-, or Myo18A siRNA-treated cells. Data represents n = 33-35 cells per condition, N = 3 independent experiments. (B) Representative images of MDA-MB-231 cells treated with NT-siRNA, βPix-siRNA or Myo18A-siRNA, stained with Sir-Actin (magenta) and Hoechst (blue). Scale bar = 20 µm. (D-E) MDA-MB-231 cells were seeded onto fibronectin-coated crossbow micropatterns 6 h prior to fixation and immunostaining. (C) Schematic of scoring system used for polarisation assessment. Cells were considered to be polarised if the centrosome was located in the front third of the cell, in front of the nucleus. Cells were considered to be unpolarised if the centrosome was in the back two thirds of the cell. (D) Representative maximum projections of cells stained with phalloidin (magenta), pericentrin (green), DAPI (blue), treated with NT-siRNA, βPix-siRNA, or Myo18A-siRNA, untransfected or transfected with GFP-βPix or GFP-βPix-Myo18A binding mutant. Scale bar = 20 µm. (E) Bar chart shows internal polarisation for conditions outlined in B as conducted based on centrosome position outlined in A. N = 3 independent experiments; n = 9 - 51 cells. One-way ANOVA; Dunnett’s multiple comparison post-test versus NT-siRNA. Error Bars: SEM.

Established front-rear cell polarity is required for cell migration (Ridley et al., 2003). We qualitatively observed rounded, well spread cells that lacked a clear front and rear following βPix/Myo18A-depletion on 2D surfaces (data not shown) and chose to further investigate cell polarity. Using the centrosome as an indicator of cell polarity (Lombardi et al., 2011). Cells were scored as “polarised” if the microtubule organising centre (MTOC) was located in the front third of the cell (from the centre of nucleus, Figure 3C). Using this method, 59% of control cells show polarisation, whereas depletion of βPix or Myo18A caused a significant reduction in polarisation, with only 40% and 31%, respectively, of cells polarised (Figure 3D, E). Having identified a role for βPix and Myo18A in internal cell polarity, we postulated that this may be through the ability of βPix ability to bind Myo18A. To test this, we undertook rescue experiments of βPix depleted cells with either full-length βPix (GFP-βPix) or a truncated βPix construct, which lacks the C-terminal PAWDENTL motif that is required for Myo18A binding (GFP-βPix ΔMyo18A {Hsu, 2014 #19)). While GFP-βPix expression in βPix-depleted cells was able to rescue cell polarisation, expression of GFP-βPix ΔMyo18A did not rescue the polarisation defect (Fig 3D, E).

In conclusion, these data show that the reduction in invasion and migration following βPix- or Myo18A-depletion is not due to a defect in matrix degradation, protrusion formation or altered RhoGTPase signalling. Rather, the functional βPix-Myo18A module regulates invasion/migration by controlling nuclear shape adaptation and maintenance of front-rear polarity.

### βPix and Myo18A are required for nuclear force transmission

Nesprins are nuclear-associated LINC complex members, which have been shown to be required for front-rear cell polarity and cell migration in constricted 3D environments (Lombardi et al., 2011; Thomas et al., 2015). The nucleus is the major obstacle to invasive cell migration in matrices with pore sizes resembling the *in vivo* ECM (Wolf et al., 2013), and is under actomyosin-mediated tension (Arsenovic et al., 2016; Woroniuk et al., 2018). We therefore hypothesised that βPix-Myo18A could potentially affect actomyosin-dependent nuclear force coupling leading to defective nuclear translocation and invasion.

We first examined whether the nucleus is under a different amount of tension when found in 3D compared to 2D environments. To examine this, we utilised a mini-Nesprin-2 FRET-based biosensor (Woroniuk et al., 2018). Here, a higher FRET readout indicates the nuclear region is under low tension, whereas a lower FRET readout indicates higher tension. Using this approach, we found MDA-MB-231 cells embedded in 3D matrices in concentrations that are permissive (0.5 mg/mL) and restrictive (2 mg/mL) to migration, were under greater tension, compared to cells plated on a 2D surface with or without matrix coating (Figure 4A, B).

**Figure 4.**
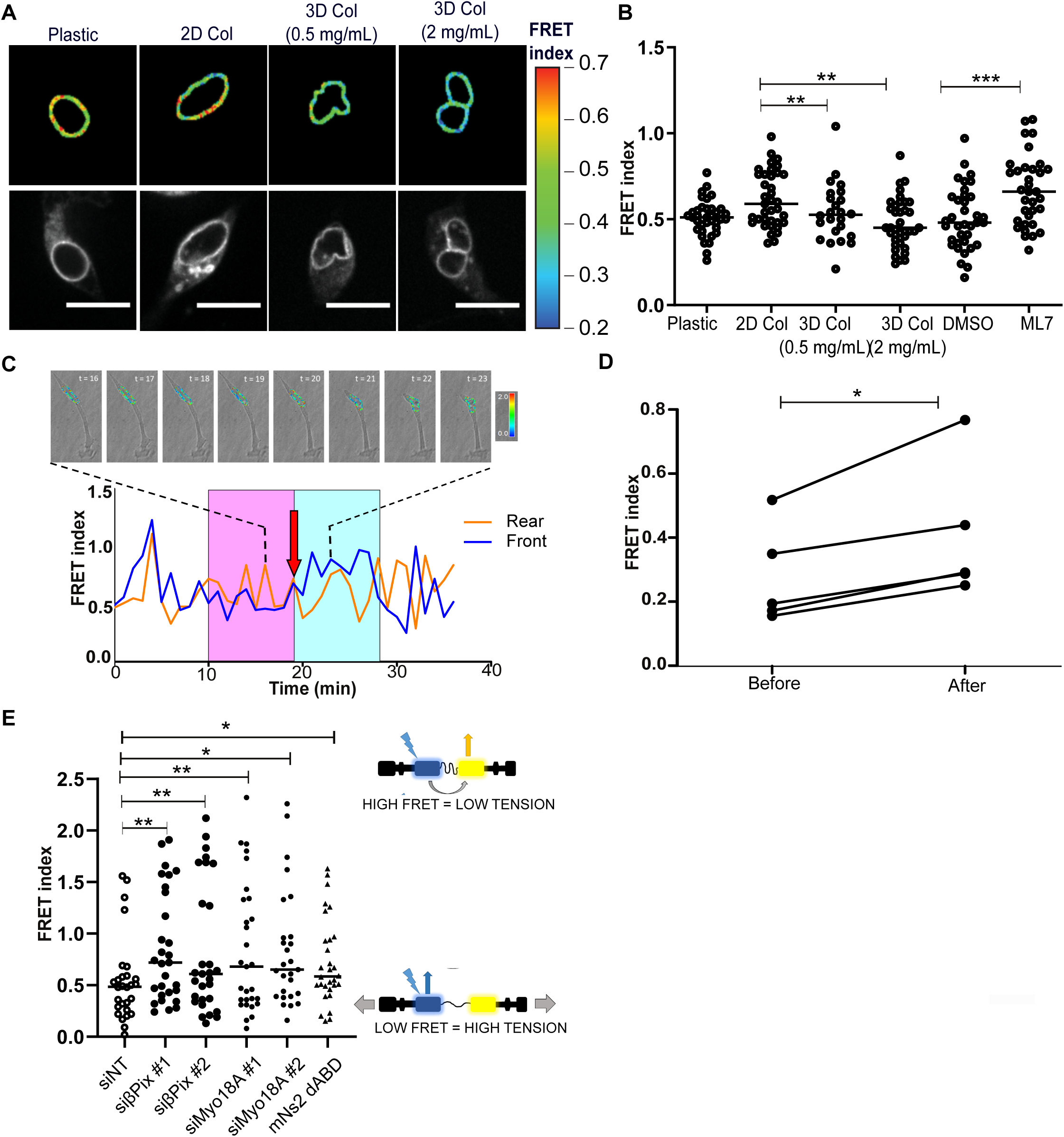
βPix and Myo18A are required for nuclear force transmission. (A) Confocal imaging of MDA-MB-231 cells expressing mN2-TS (mini-Nesprin2 tension sensor) in 2D (plastic or plastic coated with collagen) and 3D environments (0.5 mg/mL or 2mg/mL). Pseudo-coloured to represent FRET index (scale on right side). (B) The average FRET index across a central cross-section of the NE (in the horizontal plane) of MDA-MB-231 cells in differing environments (N = 4 experiments, SEM). (C) MDA-MB-231 cells expressing mN2-TS images showing FRET ratio (pseudocoloured) overlaid onto brightfield images. Images correlate to the time point range highlighted (dotted lines). Red arrow on timescale indicates moment of nuclear translocation. (D) Trace of FRET index measured for the front and rear halves of the nucleus in 2 mg/mL gel. FRET index for the ten timepoints prior to nuclear movement (pink box in C) was shown to be significantly lower than the ten timepoints after initiation of movement (blue box in C) (p = 0.067, paired t=test). (E) The average FRET index across a central cross-section of the NE (in the horizontal plane) of MDA-MB-231 cells in NT-siRNA, βPix-siRNA or Myo18A-siRNA treatment, or mini-Nesprin2 minus the actin binding domain (N = 4 experiments, SEM). Schematic of FRET sensor used shown to right, indicating low FRET equates to high tension and high FRET indicates low tension.

Next, we examined nuclear tension during restrictive 3D cell migration (cells embedded in 2 mg/mL gels). Nuclear tension of migrating MDA-MB-231 cells expressing mini-Nesprin-2 were tracked over a 40 min time period. In these conditions, our measurements show that the nucleus is under greater tension (lower FRET readout) in the front half of the nucleus before cell and nuclei movement occurs, whereas a substantial decrease in tension is observed following nuclear translocation through an ECM pore (Figure 4C-D). This was in contrast to the rear half of the nucleus, where there is no significant trend in FRET change before and after cell movement (Supplementary Figure 6A).

Having identified that there is an increase in nuclear tension in the front half of the nucleus prior to cell migration and an alteration to nuclear morphology following βPix and Myo18A loss, we examined whether βPix or Myo18A are required for nuclear force transmission. βPix- and Myo18A-depleted cells were transfected with the mini-Nesprin2 tension sensor embedded in 3D environments and imaged by live-cell microscopy. Depletion of either βPix or Myo18A resulted in a significant increase in FRET signal across nucleus (Figure 4E). A similar increase in FRET, indicating a decrease in tension, was observed following expression of a Nesprin-tension sensor which lacks the N-terminal actin binding domain (mNs2-dABD) (Davidson et al., 2020; Janota et al., 2020). Together, this data shows that force is actively applied to the front region of the nucleus during 3D cell migration. βPix and Myo18A are required for effective mechanotransduction from IACs to the nucleus in cells migrating through matrices.

### βPIX-Myo18A generates a NM2A/B gradient required for cytoskeletal-nuclear force coupling

Myo18A is known to co-assemble with NM2 (Billington et al., 2015), and NM2 isoform polarisation is required for front-rear cell polarity (Raab and Discher, 2017; Vicente-Manzanares et al., 2011). Therefore, we investigated whether the βPix-Myo18A module at adhesion sites is required for NM2 isoform mediated front-rear cell polarity and migration.

A CRISPR knock-in MDA-MB-231 cell line expressing mNeon-NM2A (named “231-mNeonNM2A” hereafter) was created to ensure endogenous levels of NM2A during live-cell analysis. Preliminary examination of 231-mNeonNM2A cells embedded in 3D matrices revealed strong accumulation of NM2A at adhesive sites (Supplementary Figure 7A). Importantly, this contrasts the distribution of NM2A in cells plated on a 2D surface, where we and others (Young and Higgs, 2018) observe less clear enrichment of NM2A at adhesive sites (Supplementary Fig. 7B). We next transfected 231-mNeonNM2A cells with mApple-Paxillin to study live-cell dynamics in 3D matrices. Similar to our fixed-cell analysis, NM2A is recruited to and enriched in adhesive sites, which also appear to resemble the sites for new protrusion assembly (Figure 5A; Supplementary Movie 1).

**Figure 5.**
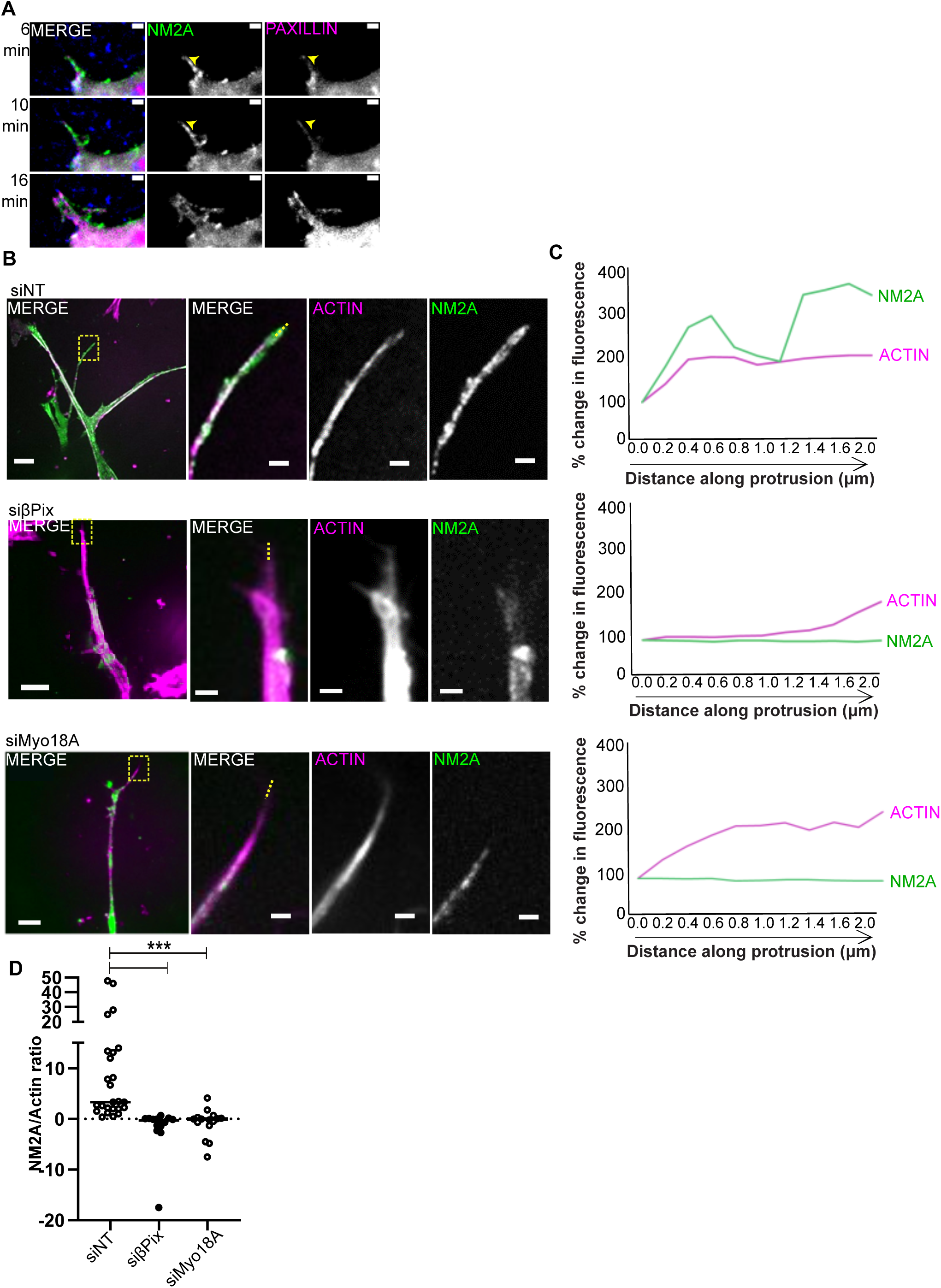
βPix and Myo18A are required for NM2A recruitment to protrusive tips. (A) MDA-MB-231 cells expressing NM2A-mNeon (green) cells transfected with mApple-Paxillin (magenta) embedded in collagen (supplemented with ∼1-2% fluorescent collagen (blue) & fibronectin gels were imaged by live-cell microscopy. Protrusive structures are indicated with yellow arrowhead. (B) Representative images of MDA-MB-231 cells expressing NM2A-mNeon (green) treated with non-targeting- (NT), βPix-, or Myo18A-siRNA embedded collagen/fibronectin gels. Cells stained with Sir-Actin (magenta). Images on left panel are expanded views, yellow boxed regions indicate zoom region shown on right. Scale bar expanded = 10 μm, zoom = 2 μm. (C) Percentage change in fluorescence signal along actin protrusion, starting at most distal point. Example fluorescence line scans of actin/NM2A from zoom images indicated by yellow dashed line shown in B. (D) Dot plot of fluorescence intensity ratio of NM2A/actin signal at 2 micron point within protrusion. n = 16-26 cells, N = 3 independent experiments. Two sample t test was used to estimate p values: *** p < 0.001.

The effect of βPix and Myo18A depletion on NM2A localisation was examined using the 231-mNeonNM2A in 3D matrices. Cells were stained with SiR-Actin to mark actin-based protrusions and imaged by live-cell microscopy. We observed no change in overall NM2A or actin intensity in pseudopodia of βPix-depleted cells compared to controls (Supplementary Fig 7C-E). However, the characteristic enrichment of NM2A fluorescence intensity towards the distal tip of actin protrusions (approx 2 μm along the length of the protrusion, toward the cell body) was absent in the βPix/Myo18A-depleted cells (Figure 5B-D). When examining cells on a 2D surface, depletion of βPix or Myo18A reduced the average intensity of NM2A staining that is in close proximity to a paxillin-based adhesion mask (Supplementary Figure 7F-G).

NM2A and NM2B form a gradient within cells plated on 2D surfaces and in 3D matrices (Shutova et al., 2017; Thomas et al., 2015). We verified these findings using MDA-MB-231 cells stained for NM2A and NM2B in 3D matrices, where NM2B accumulates in the peri-nuclear region and NM2A is distributed throughout the cell with prominent staining in pseudopodial tip regions (Figure 6A; Supplementary Figure 7A). Remarkably, depletion of either βPix or Myo18A causes a loss of the NM2A/NM2B polarisation within cells, with NM2B being less enriched in the perinuclear region and distributed throughout the cytoplasm (Figure 6A-C), whereas NM2A is no longer enriched at pseudopod tips (Figure 5B-D).

**Figure 6.**
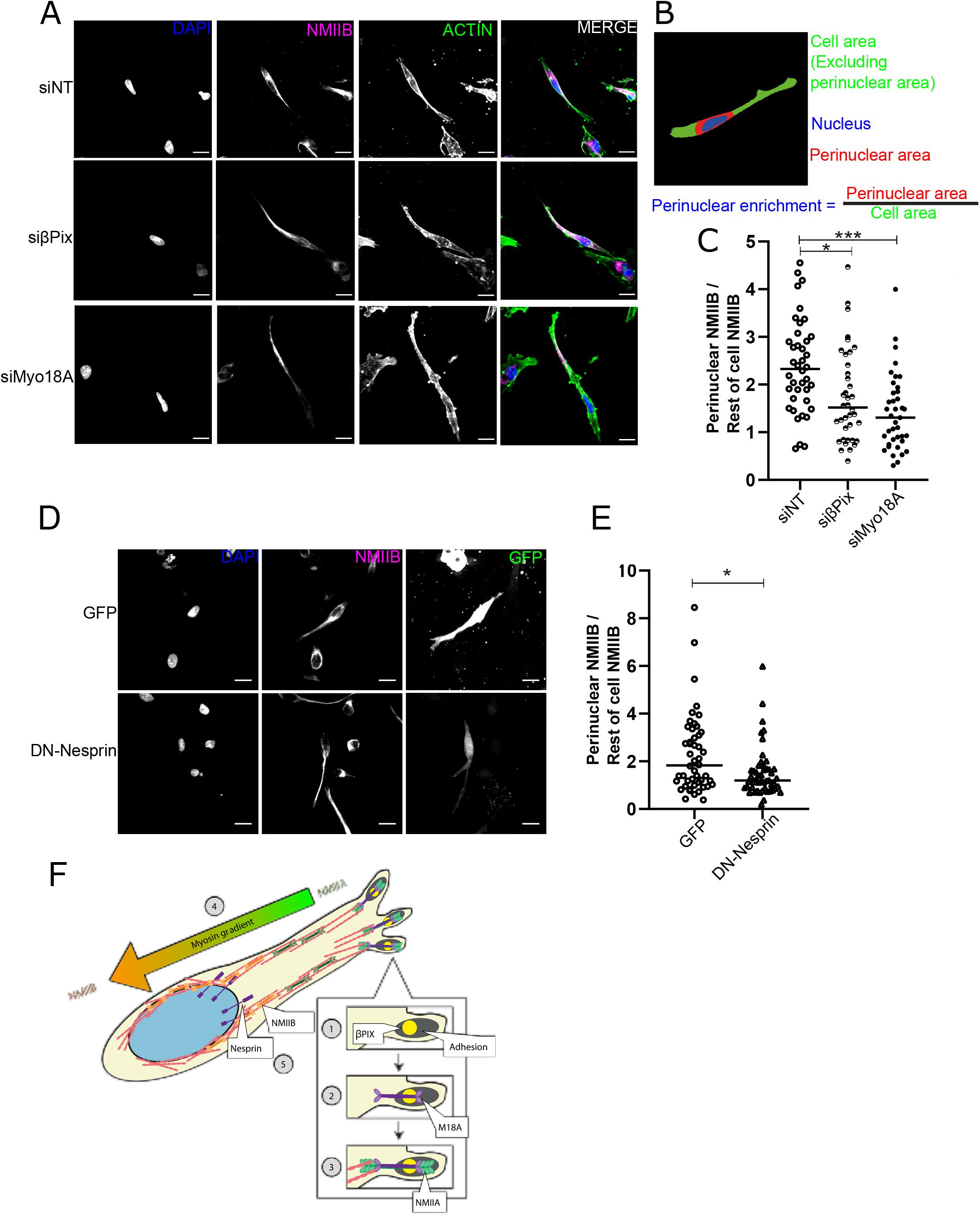
NM2A/B polarity requires nuclear force transmission mediated through βPix and Myo18A. (A) Representative images of MDA-MB-231 cells treated with non-targeting- (NT), βPix-, or Myo18A-siRNA embedded in collagen/fibronectin gels stained for DAPI (blue), NM2B (magenta), and actin (green). Scale Bar = 20 μm. (B) Schematic to show quantification method for data shown in C and E. (C) Graph shows average perinuclear NM2B intensity as a ratio of whole cell NM2B intensity (quantified as in B). Data represents n = 38-40 cells per condition, N = 4 independent experiments. (D) Representative images of MDA-MB-231 cells expressing GFP or GFP-DN-KASH (dominant negative KASH) embedded in collagen/fibronectin gels, stained for DAPI (blue) and NM2B (magenta). GFP shown in green. Scale Bar = 20 μm. (E) Graph shows average perinuclear NM2B intensity as a ratio of whole cell NM2B intensity (quantified as in B). Data represents n = 58 - 60 cells per condition, N = 5 independent experiments. (F) Model of NM2A/B gradient formation: **(1)** βPix is recruited to nascent adhesion complexes at the protruding front of the cell. **(2)** Myo18A is recruited via βPix. **(3)** Myo18A co-assembles with NM2A, the tension generated by NM2A results in the further maturation of associated adhesion complexes. **(4)** The recruitment of NM2A to the leading edge of the invading cell promotes the formation of a front-rear NM2A/B gradient. **(5)** Nesprins (-1 & -2) present on the outer nuclear membrane are able to bind to actin, and via their binding to SUN domain containing proteins present on the inner nuclear membrane, they connect the cytoskeleton and nucleoskeleton as part of the LINC complex. An actin cross linker is required for polarity maintenance and nuclear positioning. Mechanical integration of peri-nuclear actin with the nucleoskeleton is essential for the establishment of a front-rear NM2A/NM2B gradient. We propose that mechanical coupling between the nucleus and the adhesion sites is required for the establishment of this gradient and for the polarisation of the invading cell.

Having determined that expression of the Nesprin actin-binding mutant (mNs2-dABD) causes a loss of nuclear force akin to βPix/Myo18A depletion (Figure 4), and knowing Nesprin-2 is required for 3D cell migration (Thomas et al., 2015), we examined the effect of a dominant-negative KASH domain only Nesprin construct (DN-Nesprin) on NM2B polarity. DN-Nesprin expression in MDA-MB-231 cells causes a loss of NM2B polarisation (Figure 6D-E), similar to βPix/Myo18A depletion.

Together, these data show that βPix/Myo18A are required for efficient recruitment of NM2A to adhesion sites and essential for the polarised distribution of NM2A-NM2B needed for cell migration and invasion (Figure 6F). This demonstrates that migrating cells establish a connected network of actomyosin mediated force coupling from adhesion sites in protrusions to the nucleus, which underpins successful 3D invasive cell migration.

## Discussion

This study identifies the composition of core components of cell matrix adhesion sites in cells migrating within 3D matrices. Differences in the composition between adhesion complexes on stiff 2D and soft 3D matrices reveal the establishment of polarity in the actomyosin network as a novel function of adhesion sites in 3D matrices, which is required for invasive migration. 3D matrix adhesion sites are relatively enriched in βPix and Myo18A; two proteins that are needed for NM2A recruitment to adhesion sites in protrusions and NM2A/NM2B isoform gradient formation across the cell. Cells moving through 3D matrices actively translocate their nucleus, here we show dynamic nuclear force transduction between adhesion sites and the LINC complex in the nuclear membrane during 3D invasive cell migration. Loss of actomyosin connectivity at adhesion sites or the nucleus leads to a loss of nuclear force coupling and a failure to establish the NM2 isoform polarity gradient. We propose a model whereby mechanical coupling between IACs and the nucleus is required for the formation of an NM2A/NM2B gradient; the gradient in turn is essential to couple nuclear movement to cell migration (Figure 6F). We show that nuclear actomyosin force coupling from adhesion sites in 3D matrices is an essential requirement for invasive cancer cell migration.

To the best of our knowledge, this work represents the first systemic analysis of the composition of IACs in cells interacting with a 3D microenvironment. A number of novel adhesion proteins were identified here, as well as a significant proportion of established consensus adhesion components in IACs from both 2D and 3D environments (Horton et al., 2015; Horton et al., 2016). Results of the 2D IAC experiments also compared well with previously published BioID screens of paxillin (Chastney et al., 2020; Dong et al., 2016). This suggests that the core adhesion machinery and architecture remains conserved in a 3D microenvironment. The compositional changes observed in cells in a 3D matrix, compared with 2D, are highly reminiscent of those found when cellular contractility is inhibited by blebbistatin treatment, with reduced enrichment of mechanosensitive LIM-domain containing proteins and enhanced recruitment of the GIT-βPix complex (Kuo et al., 2011; Schiller et al., 2011).

A limitation of the current screen is that it is unable to ascribe which physio-chemical parameter is responsible for inducing a given change between the chosen 2D and 3D micro-environmental conditions. Substrate rigidity is likely to account for a large proportion of the changes observed, but changes such as dimensionality and ligand density between the 2D and 3D conditions may also play a role (Matellan and del Río Hernández, 2019). Future work should seek to de-convolve the contribution of each of these factors. A better understanding of the matrix parameters that are important in regulating IAC composition and signalling would allow the field to develop *in vitro* ECM models that better resemble the microenvironment found *in vivo* whilst maintaining the minimal necessary level of complexity.

Our study has identified new factors required specifically for invasive migration of breast cancer cells in 3D environments. Of the proteins enriched in 3D adhesion sites, βPix, GIT1/2 and Myo18A stood out as previously identified IAC components with unknown roles in cancer cell invasion. We show that βPIX and Myo18A are essential for invasion without affecting the ability of cells to degrade matrix or form pseudopods. We also showed that both proteins are implicated in random migration on 2D surfaces but were unable to detect changes in actin-based 3D pseudopod length, or in Rac and CDC42 activity. The latter was perhaps unexpected as βPix has previously been associated with GEF activity in fibroblasts and contains a Dbl homology (DH) domain (Kutys and Yamada, 2014; Manser et al., 1998). However, more recent publications have shown minimal GEF activity for βPix (Müller et al., 2020) and early structural studies revealed that the βPix DH domain has mutations in key residues required for activity (Aghazadeh et al., 1998). With these previous publications and the data presented here, we do not propose the lack of migration and invasion is through a cellular actin polymerisation defect in βPIX- or Myo18A-depleted cells. We were however able to confirm a previously observed cell shape change, following depletion of βPIX/M18A (Makowska et al., 2015). Cells were visibly more circular and spread than their wildtype counterparts with an inability of the cells to polarise their MTOC on defined micropatterns. This phenotype was reminiscent of cells having defects in connections of the cytoskeleton to the nucleus (Thomas et al., 2015).

Actomyosin mediated tension can be sensed by the nucleus, in particular in 3D matrix environments (Arsenovic et al., 2016; Lomakin et al., 2020; Woroniuk et al., 2018). Nuclear force transduction has been postulated to play an important role in cell migration (Davidson and Lammerding, 2014), but measurements of forces applied on the nucleus during 3D invasive cell migration have previously not been documented. Here we show that cancer cells invading into 3D collagen gels actively apply forces on the nucleus. When movement of the nucleus is stalled, despite continued protrusion formation - presumably due to the cell encountering an obstacle – tension at the front of the nucleus increases prior to nuclear movement. Following forward translocation as the nucleus moves past the obstacle, a noticeable drop of tension at the front of the nucleus is observed. The results here suggest that the nucleus is, at least in part, pulled forward by actomyosin forces applied to the nucleus via Nesprin. This idea is strengthened by a recent study which shows a specific accumulation of Nesprins at the leading edge of nuclei when crossing obstacles (Davidson et al., 2020; Davidson and Lammerding, 2014). Intriguingly, we observed pulsatile fluctuations in FRET at the front of the nucleus during 3D cell migration, which it is might be indicative of an actomyosin-dependent tugging effect. Actin cytoskeleton networks have been shown to be able to transmit forces over long ranges (Hu et al., 2017; Owen et al., 2017; Senger et al., 2019). Our results show that depletion of βPix or Myo18A leads to complete loss of nuclear force coupling, establishing that force transmission from adhesion to nucleus in 3D matrix exists and its loss might lead to an inability for cells to move in dense 3D matrices.

Why may βPix/Myo18A be required for nuclear force coupling? The actomyosin networks show a polarised distribution of NM2 isoforms that are thought to have specialised functions in the migration process (Thomas et al., 2015; Vicente-Manzanares et al., 2011); Myo18A was shown to form hetero-multimers with NM2A in cells (Billington et al., 2015). Staining of NM2A and NM2B in invasive MDA-MB231 cells in matrix revealed a pronounced gradient, with NM2B concentrated around the nucleus and NM2A enriched more towards the protrusive region. Interestingly, NM2A was particularly enriched in the protrusive tips of pseudopodia in 3D matrices, where novel adhesion sites were forming. Disruption of the βPix-Myo18A module resulted not only in loss of NM2A accumulation in the vicinity of adhesions, but also in a redistribution of NM2B throughout the cell. βPix/Myo18A-dependent recruitment of NM2A to IACs may be the initial cue that instigates a self-sorting mechanism for NM2 isoforms that results in their polarised distribution previously described for cells on 2D surfaces (Raab and Discher, 2017; Shutova et al., 2017; Vicente-Manzanares et al., 2011). This self-organising mechanism is thought to be due to the differences in NM2A/B biochemical and biophysical properties, as well as expression levels. Comparatively higher NM2A expression is supposed to lead to preferential NM2A recruitment to new actin filaments at the leading edge. NM2B has a higher affinity for ADP (higher duty ratio), and therefore remains attached to the actin filament for longer, accumulating on actin filaments over time and enriching towards the cell interior (Kovács et al., 2007; Kovács et al., 2003; Wang et al., 2003). As we do not observe any decrease in NM2A/B mRNA or protein levels following βPix /Myo18A depletion, the loss of NM2A/B polarity in our system is unlikely due to differences in expression levels. In addition, we do not identify any differences in MLC phosphorylation, thus the mis-localisation of NM2A/B does not seem to be due to a defect in regulation, although we have not analysed all potential mechanisms for NM2 activation. Instead, our data argue for a model were βPix /Myo18A are required for recruitment of NM2A to adhesion sites and give rise to the formation of an NM2 isoform gradient according to previously postulated rules of self-organisation and nuclear force coupling via the LINC complex.

NM2A can form hetero-multimers with Myo18A in cells (Billington et al., 2015). The anterior accumulation of NM2A in 3D embedded cells is known to stabilise protrusions (Rai et al., 2017) and NM2A has faster motor activity than NM2B (Barua et al., 2014) suggesting NM2A is more efficient at creating contractile forces. This leads to directional force inputs into the connected actomyosin system from the leading edge of pseudopodia and the possibility to orientate polarity. By contrast, NM2B is critical for the translocation of nuclei through restrictive pores in MDA-MB-231 cells (Thomas et al., 2015). The requirement of NM2B around the nucleus is likely due to its higher duty ratio, and the longer time NM2B spends on actin filaments, which can help generate and maintain sustained tensile force (Barua et al., 2014; Kovács et al., 2007). Recent studies have shown that actin cross linkers, such as ɑ-actinin, are required for the coherence and force connectivity of the whole actin network (Senger et al., 2019). They are also required for cell polarity as well as nuclear positioning (Senger et al., 2019). This strengthens the argument for a connected actin network driving nuclear force coupling.

## CONTACT FOR REAGENT AND RESOURCE SHARING

Further information and requests for resources and reagents should be directed to and will be fulfilled by the Lead Contact, Tobias Zech.

## EXPERIMENTAL MODEL AND SUBJECT DETAILS

### Cell culture

Human MDA-MB-231 and Human osteosarcoma U2OS cells (HTB-96™) were obtained from ATCC®. MDA-MB-231 cells were maintained in high glucose, GlutaMAX™, DMEM (Gibco, Thermo Scientific, 31966047) supplemented with 10% fetal bovine serum (Sigma-Aldrich) at 37°C and 5% CO2. Cell lines were tested at regular intervals for mycoplasma contamination using LookOut Mycoplasma PCR detection kit (Sigma-Aldrich). All cell lines were used for a maximum of 25 passages.

### Generation of stable BioID2 expressing cell lines

MDA-MB-231 cells were transfected with BioID2 using a Nucleofector™ II system (Lonza) according to manufacturer’s instructions (Program X-013, Kit V). After 2 day cells were selected with G418 (2 mg/mL, Sigma-Aldrich). Once visible colonies had formed (∼3-4 weeks) individual colonies were picked and sub-cultured. Individual colonies were then tested for expression of BioID2-construct expression via Western blotting.

### CRISPR knockin cell line generation

The CRISPR-knockin MDA-MB-231 cell line expressing mNeon-NM2A (named “231-mNeonNM2A”) was made by CRISPR-Cas9. CRISPR plasmids were transfected into MDA-MB-231 cells using TransIT®-2020 (Mirus Bio) transfection reagent in equal amounts. Clones were selected by FACS sorting and single cell cloning, then verified by immunofluorescence and western blotting.

## METHOD DETAILS

### Isolation of 3D adhesion complexes

Labelling of adhesions complexes: MDA-MB-231 cells expressing BioID-2 constructed were either seeded on to Fibronectin/Collagen coated 10cm [plastic tissue culture dishes] or embedded in 1.7 mg/mL collagen hydrogel supplemented with fibronectin. Cells were allowed to spread for 4 hours before the culture medium was supplemented with 50 μM biotin. Cells were then incubated with biotin for 16 hours prior to lysate extraction.

### Lysate extraction

2D and 3D samples were washed twice in 4°C PBS(-), followed by 4°C 50 mM EDTA PBS(-). 2D and 3D samples were then rocked at 4°C for 1 h. Collagen gels were centrifuged for 1 min at 3200 × g at 4°C and the supernatant removed. RIPA lysis buffer (150 mM NaCl, 1% (v/v) NP-40, 0.5% (w/v) Sodium deoxycholate, 0.1% Sodium dodecyl sulfate, 50 mM Tris pH 8.0, supplemented with 1x HALT protease and phosphatase inhibitor (Thermo Fisher Scientific) was then added to each sample. Samples for each condition were pooled before being repeatedly passed through a 20 mL syringe, a 1 mL syringe and a 25-gauge needle to ensure homogenisation. Samples were transferred to pre-cooled Eppendorfs and centrifuged at 14,000 × g for 15 min at 4°C. The supernatant was retained as the sample.

### Pull-down of biotinylated proteins

An equal amount of protein was used for each pulldown in every condition (diluted to an equal volume in RIPA lysis buffer). 50 mM Tris pH 7.4 was added to each sample, equal to 50% of the total volume. Pierce™ NeutrAvidin™ agarose beads (Thermo Fisher Scientific) were pre-equilibrated with 2:1 RIPA lysis buffer:50 mM Tris pH 7.4. Samples and beads were rotated at 4°C overnight. To remove each wash, samples were centrifuged at 20 × g at room temperature (RT) for 1 min before removal of the supernatant. Each sample was washed with BioID Wash Buffer #1 (2% SDS in dH2O) for 8 min on a rotator. Beads for each condition were then pooled in BioID Wash Buffer #1 and washed for 8 min. Beads were washed with 1x BioID Wash Buffer #2 (0.1% (w/v) Sodium Deoxycholate, 1% (w/v) Triton X-100, 1 mM EDTA, 500 mM NaCl, 50 mM Hepes) and 1x BioID Wash Buffer #3 (0.5% (w/v) Sodium Deoxycholate, 0.5% (w/v) NP-40, 1 mM EDTA, 250 mM LiCl, 10 mM Tris pH 7.4.) for 8 min each. Beads were then washed with 50 mM Tris pH 7.4. Proteins were eluted from the beads by the addition of a volume equal to that of the beads of 2 × reducing elution buffer. Samples were boiled for 10 min at 96°C before collection of the eluate. This was repeated a further two times. Samples were diluted 4-fold in −20°C acetone and incubated overnight at −20°C. Samples were subjected to centrifugation at 14,000 × g for 10 min at 0 °C. The supernatant was removed and protein pellets were allowed to air dry in a flow hood. Samples were re-suspended in 2x reducing sample buffer.

### Extraction of type I collagen

Approximately 12-14 frozen rat tails were thawed in 70% ethanol. The skin was removed and individual tendons were extracted, care was taken to avoid extracting the sheath. Tendons were dissolved in 0.5 M acetic acid at 4°C for 48 h. The tendon extract was centrifuged at 7500 × g for 30 min to remove debris. 10% (w/v) NaCl was added to the supernatant which was stirred for 1 h at 4°C. The extract was then centrifuged at 10,000 × g for 30 minutes. The pellet was dissolved in 0.25 M acetic acid and stirring for 24 h at 4°C. The collagen solution was dialysed (VISKING, SERA44120, MWCO 12-14 kD, regenerated cellulose) against 8 changes (twice daily) of 17.5 mM acetic acid. The dialysed collagen solution was then centrifuged at 30,000 × g for 1.5 h. The collagen solution was then stored at 4°C in a sterile container for a minimum of two weeks before use in assays. Protein concentration of the collagen was verified by both BCA Protein assay (Sigma-Aldrich, cat: BCA1) and drying via vacuum centrifugation.

### Fluorescent labelling of collagen

Rat tail type I collagen was diluted in 4°C PBS(+). Collagen was neutralized using 0.46 M sodium bicarbonate and a final collagen hydrogel concentration of 3 mg/mL was created. The gel was added to a 10 cm tissue culture dish and allowed to polymerise at RT for 1 h. The collagen gel was incubated with 50 mM Borate buffer (pH 9.0) for 10 min. 2 mg/mL of Atto-647N-NHS-ester (Sigma-Aldrich Aldrich, cat: 18373-1MG-F) was diluted in 50 mM Borate buffer (pH 9.0) and added to the collagen gel for 1 h at RT, protected from light. Labelling solution was removed from the gel, and residual ester quenched by the addition of 50 mM Tris pH 7.4 for 10 min. The gel was washed 8 × 10 minutes with PBS(+). Labelled collagen was dissolved by incubation in 0.2 M HCl at 4 °C for 4 days. Dissolved collagen was dialysed against 20 mM acetic acid overnight. When used in assays a small amount of labelled collagen (between 0.5% - 2% of total volume) was added into the prepared collagen solution, as indicated.

### Mass spectrometry sample preparation

Samples were separated by polyacrylamide gel electrophoresis. Polyacrylamide gels were stained with Instant Blue protein dye (Expedeon) for 45 min, and then washed with ddH20. Each sample was cut into 15 slices by hand using a clean glass tile and sterile scalpel. Slices were cut into cubes approximately 1 mm^3^ in size and transferred into a perforated 96-well plate (Glygen Corp). Gels pieces were de-stained with repeated 30 min incubations of 50% (v/v) Acetonitrile (ACN) / 50% 25 mM NH4HCO3 (v/v) at RT. Gel pieces were dehydrated by 2 × 5 min incubations with ACN followed by vacuum centrifugation. Peptides were then reduced via incubation with 10 mM dithiothreitol (DTT) at 56°C for 1 h. Peptides were then alkylated by incubation with 55 mM iodoacetamide (IA) for 45 min at RT whilst being protected from light. DTT and IA were removed by two rounds of washing and dehydration: 5 min incubations with 25mM NH4HCO3 followed by a 5 min incubation with ACN. ACN was removed by centrifugation, the gel pieces were then dried via vacuum centrifugation. 1.25 ng/L porcine trypsin (Promega, Cat No:V5280) in 25 mM NH4HCO3 was added to the gel pieces, which were incubated at 4°C for 45 min to allow the trypsin to permeate the gel pieces. Samples were then transferred to 37°C to allow digestion overnight. Trypsinised peptides were collected from gel pieces via centrifugation. Residual peptides were extracted with a 30 min incubation at RT with 99.8% (v/v) ACN/ 0.2% (v/v) formic acid and then 50% (v/v) ACN/ 0.1% formic acid (v/v) which were each subsequently extracted via centrifugation. The collected eluate was dried by vacuum centrifugation. Dried peptides were then stored at −20°C until resuspension prior to analysis.

Dried peptides were re-suspended in 5% (v/v) ACN in 0.1% formic acid. Each sample was separated on a Nanoacquity™ (Waters) Ultra Performance Liquid Chromatography column coupled to an LTQ-Orbitrap XL (Thermo Fisher) equipped with a nanoelectrospray source (Proxeon). MS spectra were acquired at a resolution of 30,000 and MS/MS was performed on the top 12 most intense ions

### RNA-Sequencing analysis

MDA-MB-231 cells were embedded into 1.7 mg/mL collagen supplemented with 50 µg/mL fibronectin, and total miRNA extracted using a miRNeasy kit (QIAGEN) according to the manufacturer’s protocol. RNA samples were processed and sequenced by the Centre for Genomic Research (University of Liverpool). 1g of total RNA was treated for polyA using Next Ultra Directional RNA library prep kit (New England Biosciences). Enriched RNA was amplified using the ScriptSeq v2 RNA-Seq Library Preparation protocol. After 12 cycles of amplification DNA was purified using Ampure XP beads. Denatured template DNA at a concentration of 300 pM was sequenced using two lanes of an Illumina HiSeq2500 at 2×150 bp paired-end sequencing with v1 chemistry. Basecalling and de-multiplexing of indexed reads was performed by CASAVA version 1.8.2 (Illumina. Sequences were trimmed to remove Illumina adapter sequences using Cutadapt version 1.2.1, any reads which matched the adapter sequence over at least 3 bp were trimmed. Low quality bases were removed using Sickle version 1.200 with a minimum window quality score of 20. Finally after trimming all reads shorter than 10 bp were removed.

Quantification of expression of transcripts from the trimmed datasets was performed using Salmon v0.12.0 (Patro et al., 2017), transcripts were mapped against Homo Sapiens transcriptome (Version 86, EMBL). Gene level count matrices were generated and differentially expressed genes were determined using the DESeq2 package (Love et al., 2014; Soneson et al., 2015).

### DNA transfections

Cells to be transfected were seeded at 2 × 10^5^ cells per well in a 6-well plate ∼ 16-20 h prior to transfection. MDA-MB-231 cell transfections were performed using TransIT®-2020 transfection reagent or Lullaby transfection reagent (OZ Bioscience).; 1 μg of each plasmid was used for all transfections (2 μg total). Cells were transfected for 24 h before imaging. Before imaging, for 2D cell culture, cells were plated onto glass bottom 35 mm dishes (SPL Confocal 35-mm clear coverglass-bottom petri dish, #101350) pre-coated for 1 h (4 degrees) with 50 μg/mL collagen (as prepared below) and 10 μg/mL fibronectin (Sigma-Aldrich, FL1141). For 3D cell culture, cells were embedded in matrices 8-18 h before imaging.

#### Plasmids used in this study include

pcDNA3.1 mycBioID (a gift from Kyle Roux (Addgene plasmid # 35700; http://n2t.net/addgene:35700; RRID:Addgene_35700), myc-BioID2-MCS (a gift from Kyle Roux (Addgene plasmid # 74223; http://n2t.net/addgene:74223; RRID:Addgene_74223), MCS-13X-Linker-BioID2-HA (a gift from Kyle Roux (Addgene plasmid # 80899; http://n2t.net/addgene:80899; RRID:Addgene_80899), mApple-Paxillin-22 (Addgene, #54935), mN2-TS (Woroniuk et al. 2018) Raichu-Rac1 (gift from Patrick Caswell (Itoh et al., 2002)), Raichu-Cdc42 (gift from Patrick Caswell, original (Itoh et al., 2002)), Rac- and CDC42-FLARE.dc1g (gifts from Klaus Hahn, University of North Carolina at Chapel Hill), GFP-Myosin18Aa (Gift from Martin Humphries, University of Manchester), mCherry-Talin-1 (Gift from David Critchley, University of Leicester), GFP-Paxillin (Gift from Patrick Caswell, University of Manchester), mCherry-Vinculin (Gift from Christoph Ballestrem, University of Manchester), dominant-negative (tm) Nesprin and mini Nesprin dABD (Lombardi et al. 2011; Gift from Akis Karakesioglou, Durham University), GFP-βPIX, GFP-βPIX Δ M18A (Zech Lab, this study; (Hsu et al., 2014)). CRISPR plasmids including all-in-one CRISPR-Cas9 vector, MS2-CtIP and donor vector, PITCh-mNeonGreen-guideMYH9 were designed according to (Sakuma et al., 2016; Nakade et al., 2018)

#### Plasmids generated in this study

myc-BioID2-Paxillin was generated by the amplification of Paxillin via PCR from GFP-Paxillin using the primers TCA TGC GAA TTC ATG GAC GAC CTC GAC GCC and TGA GAC AAG CTT CTA GCA GAA GAG CTT GAG. The PCR product and myc-BioID2-MCS were digested with EcoR1 and HindIII, purified and ligated.

A start codon and kozack sequence were introduced into MCS-13X-Linker-BioID2-HA via mutagenesis (Forward: CCC GCC TCC ACC GGA TCC CAT GGT GGA AAC ACC ATG GGA TCC GAA TTC GAA TC, Reverse: GAT TCG AAT TCG GAT CCC ATG GTG TTT CCA CCA TGG GAT CCG GTG GAG GGC GG) to produce the ATG-13X-Linker-BioID2-HA vector.

To generate the Talin-13X-Linker-BioID2-HA, Talin was amplified from mCherry-Talin-1 (Forward: GAT CCA GAA TTC CAC CAT GGT TGC GCT TTC GCT G, Reverse: TGG ATC GAA TTC GTG CTC GTC TCG AAG CTC). The PCR product and MCS-13X-Linker-BioID2-HA were digested with EcoRI, purified and ligated.

The all-in-one CRISPR-Cas9 vector was generated to express Cas9 and MS2-containing sgRNAs for donor cleavage and genome targeting in accordance with the previous report (Nakade et al., 2018). The inserted oligonucleotides for genome targeting were as follows: sense, CAC CGT GCA GGA GAA GAG GCT TAT T and antisense, AAA CAA TAA GCC TCT TCT CCT GCA C. The PITCh-mNeonGreen-guideMYH9 was generated by PCR and In- Fusion cloning method (Takara) to insert mNeonGreen at the 3’ end of MYH9 coding sequence with 40-bp microhomologies in accordance with the previous reports (Sakuma et al., 2016; Nakade et al., 2018).

### 3D cell embedding

For 3D cell embedding, gels were prepared fresh on the day of use. All solutions and collagen were kept at 4 °C during preparation of the gel. The required amount of rat tail type I collagen was diluted with sterile ddH20 in order to achieve the desired final gel concentration. 10x DMEM equal to 10% of the final volume was added to the collagen gel. The pH of the gel was neutralised via the addition of 0.46 M Sodium Bicarbonate equal to 10 % of the final volume. Gels were supplemented with fibronectin (Sigma-Aldrich) to a final concentration of 10 µg/mL. The desired number of cells were trypsinised and pelleted via centrifugation. Cells were suspended in collagen/fibronectin gel solution by gentle pipetting. Cell suspensions were quickly transferred either to a cell imaging dish (SPL 35-mm dishes) or 6 well plate depending on the assay. Collagen gels were then transferred to a humidified CO_2_ incubator at 37°C to allow gel polymerisation. After 1 h, full growth medium was added carefully to the top of the gels and cells were returned to 5% CO_2_/37°C until imaging/fixation.

### siRNA treatment

MDA-MB-231 cells: 10 nM of each siRNA oligonucleotide was introduced using Lullaby® transfection reagent (Oz Biosciences) following manufacturer’s instructions. U2OS cells: 50 nM of each siRNA oligonucleotide was introduced using Lipofectamine ® RNAiMAX (Thermo Fisher Scientific) following manufacturer’s instructions.

Oligonucleotides for siRNA included: human βPix (synthesized by Qiagen), human myosin-18A (synthesized by Qiagen)

βPix siRNA – #1 - AACAATCAACTGGTAGTAAGA (Qiagen S104239011), #2 CAAGCGCAAACCTGAACGGAA (Qiagen S104308997)

Myosin-18A siRNA – #1 CACGAACTGGAGATGGATCTA (Qiagen SI04273668), #2 CAGTCGTGTCAGAAGAAGTTA (Qiagen S104318034)

For all knockdown experiments, cells were subjected to siRNA treatment at 24, 72 h post-seeding. Where one siRNA treatment is indicated this is a pooled siRNA mix of the siRNAs listed above. Cells were analysed 96 - 120 h post-original transfection. All knockdown experiments were undertaken at least three independent times unless otherwise stated.

### Immunofluorescence staining

For cells plated on 2D surfaces, at appropriate time points following cell plating, cells were fixed by removal of medium and addition of 4% formaldehyde (Electron Microscopy Sciences, Hatfield, PA) in PBS for 20 min at RT, washed 3x with PBS, permeabilized with 0.25% Triton X-100 in PBS for 15 min, washed 3x with PBS, and incubated in blocking buffer (10% bovine serum albumin (BS)] Sigma-Aldrich, A7906 in PBS) for 30 min. For cells embedded in 3D surfaces, cells were fixed by removal of medium and addition of 4% formaldehyde in PBS for 30 min at RT, washed 3x with PBS, permeabilized with 0.25% Triton X-100 in PBS for 30 min, washed 3x with PBS, and incubated in blocking buffer for 1 h.

All cells were incubated with primary antibody (in PBS plus 10% blocking buffer) for 1.5 h at RT, followed by 3 × PBS washes, and incubation in secondary antibody and Acti-Stain 670 Phalloidin (Cytoskeleton, inc), plus Hoechst (Thermo Scientific, H3570) or DAPI (Thermo Scientific, D1306) in PBS plus 10% blocking buffer for 1.5h at RT. Cells were washed in 3 × PBS, and stored at 4°C in PBS before imaging by confocal microscopy.

#### Primary antibodies

Mouse monoclonal anti-Paxillin (1/50, BD biosciences #610569), Polyclonal rabbit Anti-βPix (Millipore 1:200, 07-1450-l), Rabbit polyclonal anti-myosin Heavy Chain 9 (1:100 Cell Signalling Solution Technology #3403), Mouse monoclonal anti-myosin Heavy chain 10 (3H2) (1:200 Abcam ab684), Rabbit Polyclonal anti-pericentrin (1:2000, Abcam ab4448), Mouse monoclonal YAP antibody (1:1000, Santa Cruz Biotechnology sc-101199). Rat monoclonal anti-α5β1-integrin (1:200, Mab11, homemade Morgan lab).

#### Secondary antibodies

Alexa-Fluor® 488, 594 & 647 anti-mouse, rabbit & rat IgG antibodies (Jackson ImmunoResearch). All secondary antibody stock solutions were diluted to a concentration of 1.7 mg/mL and used at a dilution of 1:200. Alexa-Fluor®-647 or Texas Red® conjugated Phalloidin (Molecular Probes, Invitrogen) was diluted to approximately 6.6 µM in methanol and used at a dilution of 1:400.

### Transwell inverted invasion assay

Matrigel (Corning) was diluted to 5 mg/mL with PBS (-) and supplemented with fibronectin to a final concentration of 25 g/mL. The Matrigel mixtures were polymerised in transwell inserts (Corning) at 37°C for 1 h. Inserts were then inverted and 8 × 10^4 cells were seeded onto the transwell filter. Cells were allowed to adhere 3-6 h before inversion of the insert. Serum-free culture medium was added to the wells of the transwells plate while culture medium containing 10% FBS and 20 ng/mL EGF (Sigma) was added to the top of the matrigel. After 5 days of culture, samples were fixed with 4% PFA for 30 min and then permeabilized by treatment with 0.1% Triton X-100 in PBS(-) for an additional 30 min. Samples were stained with DAPI and stacks imaged every 10 μm. For negative invasion controls, GM6001 (MT1-MMP inhibitor) treatment was used at 5 μM GM6001 was added for the duration of the experiment.

### Rac/Cdc42 Pull-down assays

Cdc42 and Rac1 Pull-Down assays were carried out using Cdc42/Rac1 Activation Assay Biochem Kits (Cytoskeleton, inc), following manufacturer’s instructions. Briefly, cells were washed in ice-cold PBS and lysed in ice-cold lysis buffer supplemented with protease inhibitor cocktail following 96 h of siRNA treatment. Approximately 800 μg protein were loaded onto PAK-PBD beads and subjected to pull-down assay protocol. Samples were analysed by SDS--page and western blot analysis.

### Western blot analysis

For detection of siRNA depletion: Cells were extracted 96 - 120 h post-transfection. Cells were washed 2× in ice-cold PBS and lysed in radioimmunoprecipitation assay (RIPA) buffer:0 mM Tris-HCl, 150 mM NaCl, 1% Triton X-100 (w/v), 0.1% SDS (wt/v), and 1% sodium deoxycholate (w/v) supplemented with Halt protease and phosphatase inhibitors (Thermo Fisher Scientific) by rocking at 4°C for 10 min. Just before SDS-PAGE, the protein sample was mixed 1:5 with 5x Sample Buffer (15% SDS, 312.5 mM Tris pH 6.8, 50% Glycerol, 16% β-Mercaptoethanol). Proteins were separated by 12-20 % SDS–PAGE and transferred to a polyvinylidene fluoride membrane (Millipore, Billerica, MA). The membrane was blocked with TBS-T (20 mM Tris-HCl, pH 7.6, 136 mM NaCl, and 0.1% Tween-20) containing 3 % BSA for 1 h, then incubated with primary antibody at 4°C overnight. After washing with TBS-T, the membrane was incubated with HRP-conjugated secondary antibody (Bio-Rad, Hercules, CA) for 2 hat RT. Protein signal was measured by fluorescence-based detection or chemiluminescence. For fluorescence based detection, membranes were imaged using a LICOR® ODYSSEY® Sa infrared imaging system. For chemiluminescence based detection, membranes were developed using Clarity Western ECL Substrate (BioRad, 1705061) according to manufacturer’s instructions and imaged using a Chemidoc™ Touch (BioRad) imaging system.

#### Primary antibodies

Polyclonal rabbit anti-βPix (Millipore 1:1000, 07-1450-l), Monoclonal mouse anti-Myosin18A (1:200, Santa Cruz, H-10, 365328), Monoclonal mouse anti-Cdc42 monoclonal antibody (1:250, Cytoskeleton, Cat. # ACD03), anti-Rac1 (1:250, Cytoskeleton, Cat. # ACC03, Polyclonal rabbit anti-GAPDH (1:10,000: AB2302; EMD Millipore), Monoclonal mouse anti-phospho-myosin-light-chain (1:500, Cell Signalling #3675), myosin-light-chain (1/500, Cell Signalling #8505S).

For the detection of biotin, membranes were incubated with Alexa Fluor®-680 conjugated streptavidin (1.8 mg/mL, Jackson ImmunoResearch) diluted 1:2000 in 5 % BSA (in TBST). Membrane blots were incubated with either Alexa Fluor® 680 or 790 conjugated (Molecular Probes, Invitrogen) or HRP conjugated (Pierce) secondary antibody diluted 1:10000 in blocking buffer.

### Gelatin degradation assay

The QCM™ gelatin invadopodia assays were performed according to manufacturer instructions (Millipore). 100,000 MDA-MB-231 cells treated with the indicated siRNA were plated per well of a 24-well plate. Cells were fixed and stained for actin and DAPI following 24 h incubation. The total number of degradation spots formed per cell and percent of cells associated with at least one degradation spot was quantified. Only cells with at least one degradation spot were analysed.

### Wound healing assay

U2OS cells were treated with siRNA as indicated. Cells were then cultured in monolayers were cultured in 12-well plates (3×10^5^ cells per well) with culture-Insert 2 Well in µ-Dish 35 mm (ibid) for 24 h before the insert was removed. Phase-contrast images were acquired with a Zeiss Widefield system equipped with 20xNA lens. Images were acquired every 30 min for a 40 h period. Wound area was determined using ImageJ software.

### Internal polarity assay

Coverslips with micropatterned crossbow shapes coated with fluorescent fibronectin (excitation at 650 nm) were obtained from CYTOO SA (USA). CYTOOchips™ were placed in a 6-well tissue culture plate and treated with culture medium before addition of cells. MDA-MB-231 cells treated with siRNA as stated, were trypsinized and diluted to a concentration of 15,000 cells per mL. Cells were added to each well, and left undisturbed for 15 min, before being moved to the culture incubator for 1 h. Unattached cells were removed by multiple rounds of careful PBS(-) washing. Cells were then returned to a tissue culture incubator allowed to spread 6 h prior to fixation.

### Imaging details

Unless otherwise stated, images were recorded with a Marianas spinning disk confocal microscope (3i) using a 63x 1.4 NA or 10x 0.45 NA Zeiss Plan Apochromat lens and either an Evolve electron-multiplying charge-coupled device (Photometrics) or FLASH4 sCMOS (Hamamatsu) camera.

### TIRF imaging

Imaging of cells was carried out on the Marianas spinning disk confocal microscope with an attached motorised TIRF module (Zeiss) and αPlan-Apochromat 100x/1.46 oil (Zeiss) TIRF objective was used. Images were acquired using a Hamamatsu ORCA Flash 4.0 CMOS camera with 2x binning. For imaging of fixed samples via TIRF microscopy cells were seeded onto glass-bottomed cell imaging dishes (SPL 35-mm dishes). Samples were then prepared according to the protocol described above for immunofluorescence staining. With the exception that dishes were left in a final PBS wash (rather than mounting medium) and imaged immediately.

### FRET Imaging

Imaging of cells for determination of FRET was carried out on the Marianas spinning disk confocal microscope, for each image, five channels were captured; one channel detecting a 447-517 nm range under a 445 nm excitement with a 1000 ms exposure (denoted ‘Donor’ channel), another detecting a 515-569 nm range under 514 nm excitement with a 200 ms exposure (denoted ‘Acceptor’ channel), a third detecting a 515-569nm range under 445 nm excitement with a 1000 ms exposure (denoted ‘Transfer’ channel), and a further two channels consisting of a brightfield illumination of 100ms and a far-red channel (652-732 nm range under 647 nm excitement with an exposure of 1000 ms) for imaging fluorescent collagen. For singular images, the laser power was set to 100%; for time-lapse images, the power was set to 20%. Images were taken once every min for time-lapse images.

## QUANTIFICATION AND STATISTICAL ANALYSIS

### Mass spectrometry data analysis

Obtained spectra were searched against a database containing both manually reviewed (UniProt) and unreviewed (Trembl) proteins (Uniprot-Consortium, 2019) using Mascot Sever (version 2.3.2, Matrix Science) (Perkins et al., 1999). Mascot search results were imported into Scaffold (Version 4) for statistical analysis. Results were also searched against X!Tandem to increase the confidence of protein identification. Protein and peptide confidence thresholds were set at 99% and 95% respectively.

Proteins were considered to be enriched in a given condition if they differed statistically significantly from the appropriate BioID2 only control (Fisher Exact, P < 0.05, Benjamini–Hochberg corrected). Proteins assigned to unreviewed Uniprot entries within the enriched protein list were then manually curated to identify if an appropriate reviewed entry exists. Proteins were reassigned to a reviewed entry if the sequence of an unreviewed entry was contained wholly within that of a reviewed entry or if the identified peptides were all contained within a reviewed entry. Entries which are contained in more than 33% of CRAPome controls were also excluded from the enriched list (Mellacheruvu et al., 2013)) in addition endogenously biotinylated enzymes involved in metabolic carboxylation and decarboxylation reactions were removed from protein lists (Cronan, 1990). Enriched protein lists were manually curated on the basis of their UniProt annotation for subcellular localisation to remove extracellular matrix/region proteins. Enriched protein lists were also manually curated to ensure that duplicate entries were not present, for display purposes the total spectra counts were combined for the entries P14923 and B4DE59 (junction plankoglobin, JUP).

Visualisation of data and protein interaction networks was conducted using Cytoscape (version 3.7.1). Interaction networks were obtained from the STRING 10.5 database, interactions with a confidence score > 0.4 on the basis of experimental and database evidence were included in the interaction network.

### Invasion assay (Cell Invasion Index)

To determine invasiveness of MDA-MB-231 cells (treated as indicated), first the integrated density of each optical section (DAPI stain taken every 10 μm) was calculated using ImageJ. To calculate an invasion index score the sum of the integrated density of 40 μm into the gel was divided by the negative sum of all invasive optical slices which was then normalised against the siNT control. N = ≥ 4 independent experiments.

### Wound healing analysis

Wound closure of U2OS cells (treated as indicated) was examined every 30 min for a 40 h period. Area of wound closure in every image acquired over 40 h period, using ImageJ software. Data shown represents percentage of wound closure compared to time-point zero. Data represents N = 8-10 wound area per condition, N = 2 independent experiments.

### Gelatin degradation

MDA-MB-231 cells (treated as indicated) were incubated for 24 h before fixing and staining with phalloidin and DAPI. Cells were imaged using spinning disk confocal microscopy. Following image acquisition cells were analysed using ImageJ software. Two parameters were examined: 1. Percentage of cells in randomly (Roulette) selected ROI associated with one or more gelatin degradation spots, and 2. Number of degradation spots per cell. Data represents n = 157-217 cells per condition, N = 2 independent experiments.

### Actin protrusion length

MDA-MB-231 cells (treated as indicated) were embedded in 3D matrices before staining with 200 nM SIR-Actin (Cytoskeleton, Inc) for 2 h prior to imaging. Cells were imaged using spinning disk confocal microscopy. Following acquisition, length of actin protrusions were examined using ImageJ software, measuring the full length of actin protrusions from tip to cell body. n = 35-40 cells per condition, N = 3 independent experiments.

### Nuclear circularity

Nuclear circularity of MDA-MB-231 cells (treated as indicated) was calculated using ImageJ software measuring the longest versus shortest axis of the nucleus in the direction of protrusion formation.

### FRET data Analysis

Analysis of raw images was conducted using the SlideBook 6 software (Intelligent Imaging Innovations, Inc) including the FRET module. Bleedthrough values for mTFP1 were determined using the ‘Compute FRET Bleedthrough’ functionality of the FRET module, imputing images of the mN2-TFP control construct taken under the same conditions as the mN2-TS images. To analyse FRET readings at the nuclear envelope, and exclude non-specific readings, a mask was manually drawn over the NE signal (as visualised by the Acceptor channel). All measurements and bleedthrough calculations were then conducted using pixels within the mask, subtracting for background intensity that was calculated from a region outside of the cell that showed no specific fluorescent signal.

### Presence of NM2A in protrusions

231-mNeonNM2A cells (treated as indicated) were embedded in 3D matrices before staining with 200 nm SIR-Actin (Cytoskeleton, Inc) for 2 h prior to imaging. Cells were imaged using spinning disk confocal microscopy. Following acquisition, NM2A and actin intensity was examined using Image J software following protocol outlined (www.unige.ch/medecine/bioimaging/files/1914/1208/6000/Quantification.pdf). NM2A in whole actin protrusion intensity was measured for “whole protrusion” quantification. Data shown represents actin intensity alone, NM2A intensity alone, and NM2A:actin ratio. N = 23-30 cells. NM2A within 2 μm of most distal region of protrusion tip. Data shown represents NM2A:actin ratio at 2 μm within protrusion tip. n = 16-26 cells.

### Perinuclear NM2B

MDA-MB-231 cells were embedded within 1.7 mg/mL collagen hydrogels supplemented with 50 μg/mL fibronectin and cultured for 16 hours prior to fixation and staining. Stained collagen gels were imaged on a 3i spinning disk confocal microscope, acquiring 40 slices 0.5 μm apart. Slices were background subtracted before collapsing as a SUM intensity profile. A nuclear mask was generated from the DAPI stain, while a whole cell mask was generated from the phalloidin stain. The perinuclear region was defined as a 5 μm band around the nucleus (N = 5, n = 8-10).

## Author contributions

Conceptualization: D.N, L.Y, T.W, L.M, M.M and T.Z

Methodology: D.N, L.Y, T.S, T.Y, M.M and T.Z

Software: D.N, L.Y, T.W

Validation: D.N, L.Y, T.W, K.W, A.C.O, E.M, L.B, and T.Z.;

Formal analysis: D.N, L.Y, T.W, L.B, E.M, M.M and T.Z

Investigation: D.N, L.Y, T.W, K.W, A.C.O, E.M, P.C, L.B, T.S, T.Y. and T.Z.;

Resources: L.M, M.M, P.C, T.Y and T.Z;

Writing – Original Draft: D.N, LY, T.W and T.Z;

Data curation: D.N, L.Y, T.W, M.M and T.Z,

Visualization: D.N, L.Y, and T.Z,

Supervision, Project and Funding Acquisition: L.M, T.Y, M.M and TZ

## ACKNOWLEDGMENTS

We thank the Liverpool Biomedical Imaging Facility, the Centre for Cellular Imaging (BBSRC grant BB/M012441/1) and the Centre for Genomic Research for their support and access to equipment. The mNeonGreen cDNA was used under a license agreement with Allele Biotechnology and Pharmaceuticals, Inc. L.M. thanks Cancer Research UK for core funding. P.C. was funded by Wellcome Trust Centre for Cell-Matrix Research by grant 203128/Z/16/Z. T.Y. and T.S. were funded by the Japan Society for the Promotion of Science (17H01409). D.N. was funded by Wellcome Trust PhD studentships (105350/Z/14/ A). L.Y. is funded by MRC project grant MR/S008632/1 to T.Z. T.W. is funded by the Discovery Medicine North Biotechnology and Biological Sciences Research Council doctoral training program (BB/M011186/1/1797330). E.M. was funded by a North West Cancer Research grant (1032) to TZ. L.B. was funded by a Breast Cancer Now grant (2014MayPR292) to TZ.

The authors declare no competing financial interests.

---

**Supplementary Figure 1.**
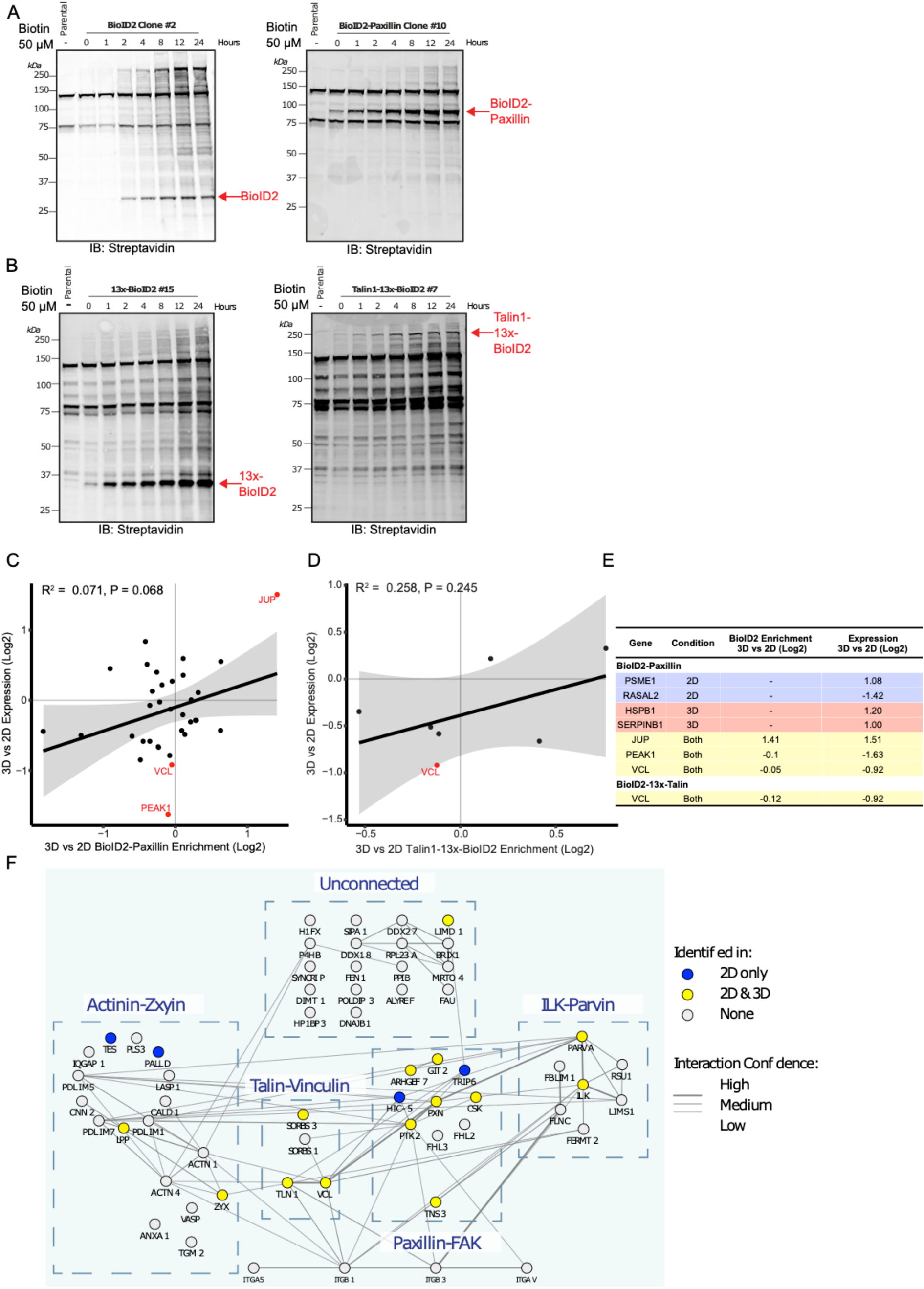
Comparison of IAC-associated proteins in MDA-MB-231 cells cultured in a 2D vs 3D microenvironment. (A, B) Biotinlyation time-course with various BioID2-construct expression. Biotin incubation 50 μM for the times indicated. (C) Expression changes identified in MDA-MB-231 cells cultured in 2D vs 3D by RNA-seq. Scatterplots showing 3D vs 2D gene expression as assessed by RNAseq vs. 3D vs 2D protein enrichment in (C) BioID2-Paxillin (D) Talin1-13x-BioID2 datasets. Red points represent genes with expression changes that are statistically significant (P < 0.05, Benjamini-Hochberg corrected). (E) Table showing genes with statistically significant expression changes 3D vs 2D that were identified in either the BioID2-Paxillin or Talin1-13x-BioID2 datasets. (F) Diagram showing coverage of the ‘consensus’ adhesome from (Horton et al., 2015) by the BioID2-Paxillin & Talin-13x-BioID2 datasets. Blue dashed boxes enclose the 4 signalling axes described by Horton *et al.,* Left to right: actinin-zyxin, Talin-Vinculin, Paxillin-FAK, ILK-kindlin; top: unconnected ‘consensus’ adhesome proteins. Interaction network obtained from STRING 10.5 (Experimental and Database interactions only, confidence > 0.4). Figure corresponds to Figure 1.

**Supplementary Figure 2.**
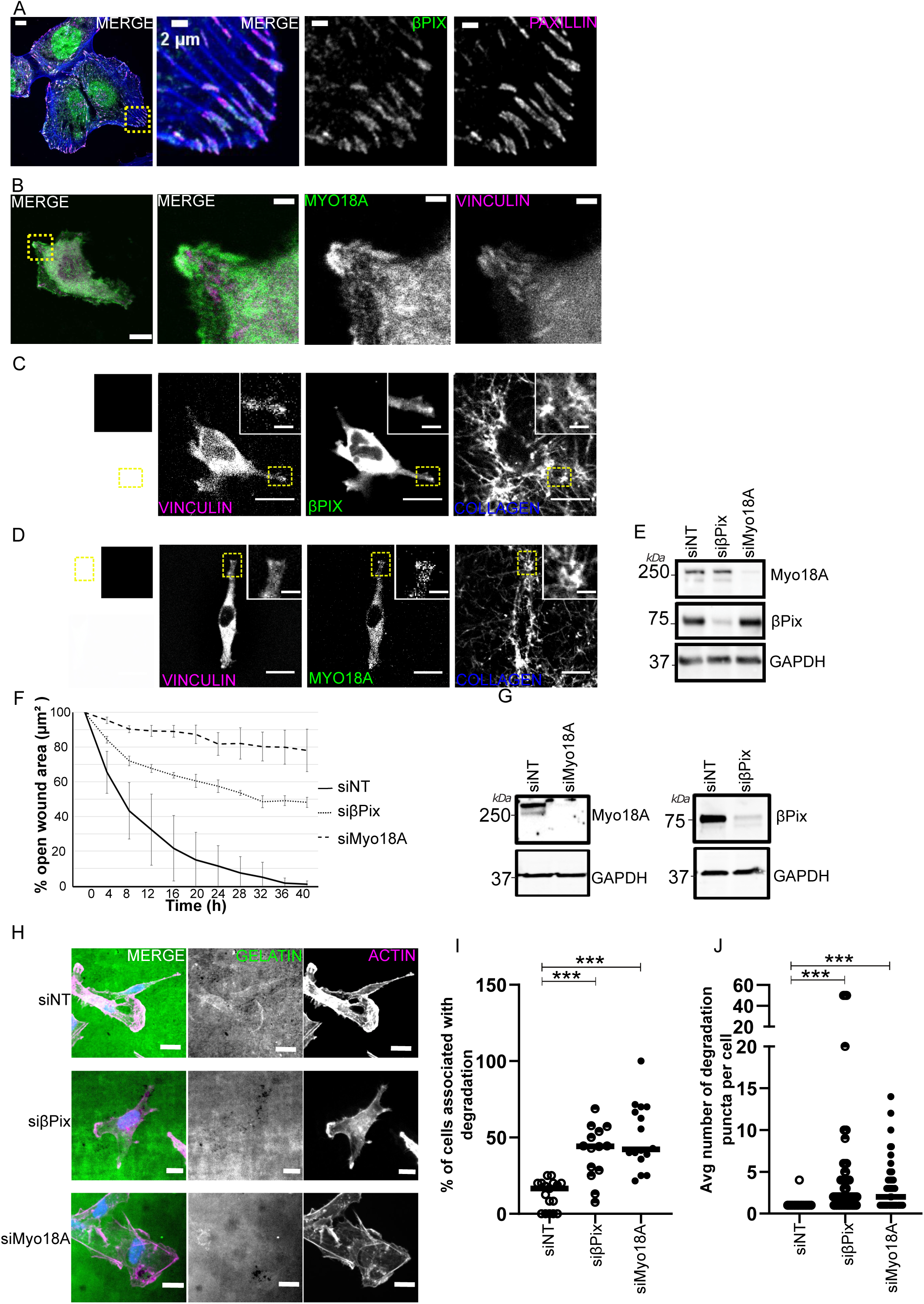
βPix and Myo18A localise to adhesive sites and are required for cell invasion. (A) βPix enriches in IACs in U2OS cells. Representative image of U2OS cells fixed and stained with phalloidin (blue), Paxillin (magenta), and βPix (green). Boxed region in expanded view indicates region of magnified view. Images are single Z slice projections. Scale bar in expanded view = 10 µm, in magnified view = 2 µm. (B) Representative image of MDA-MB-231 cells transfected with mCherry-Vinculin (magenta) and GFP-M18A (green) plated on fibronectin/collagen-coated coverslips. Boxed region in expanded view indicates region of magnified view. Images are single Z slice projections. Scale bar in expanded view = 10 µm, in magnified view = 2 µm. (C) Representative image of MDA-MB-231 cells transfected with mCherry-Vinculin (magenta) and GFP-βPix (green) embedded in collagen (supplemented with ∼1-2% fluorescent collagen (blue) & fibronectin gel). Images are single Z slice projections. Scale bar in expanded view = 20 µm, in magnified view = 2 µm. (D) Representative image of MDA-MB-231 cells transfected with mCherry-Vinculin (magenta) and GFP-Myo18A (green) embedded in collagen (supplemented with ∼1-2% fluorescent collagen (blue) & fibronectin gel). Images are single Z slice projections. Scale bar in expanded view = 20 µm, in magnified view = 2 µm. (E) Representative Western blot of MDA-MB-231 cells treated with non-targeting-(NT), βPix-, or Myo18A-siRNA. Western blot probed for Myo18A, βPix-, and GAPDH (loading control). (F) U2OS cells treated with non-targeting-(NT), βPix-, or Myo18A siRNA were plated on 2D surfaces and underwent wound healing. Graph shows wound area measured 0 h - 40 h post wound. Data represents n = 8-10 wound area per condition, N = 2 independent experiments. (G) Representative Western blot of U2OS cells treated with non-targeting- (NT), βPix-, or Myo18A-siRNA. Western blot probed for Myo18A, βPix-, and GAPDH (loading control). (H) MDA-MB-231 cells treated with non-targeting- (NT), βPix-, or Myo18A-siRNA and plated on FITC-labelled gelatin (green). Cells were fixed and stained for actin (magenta) and DAPI (blue) after 24 h of seeding. Scale bar in expanded view = 10 µm. (I) Quantification of percentage of cells associated with one or more degradation spots under a cell. (J) Quantification of average number of degradation areas formed per cell. Data represents n = 157-217 cells per condition, N = 2 independent experiments. Two sample t test was used to estimate p values: *** p < 0.001. Figure corresponds to Figure 2.

**Supplementary Figure 3.**
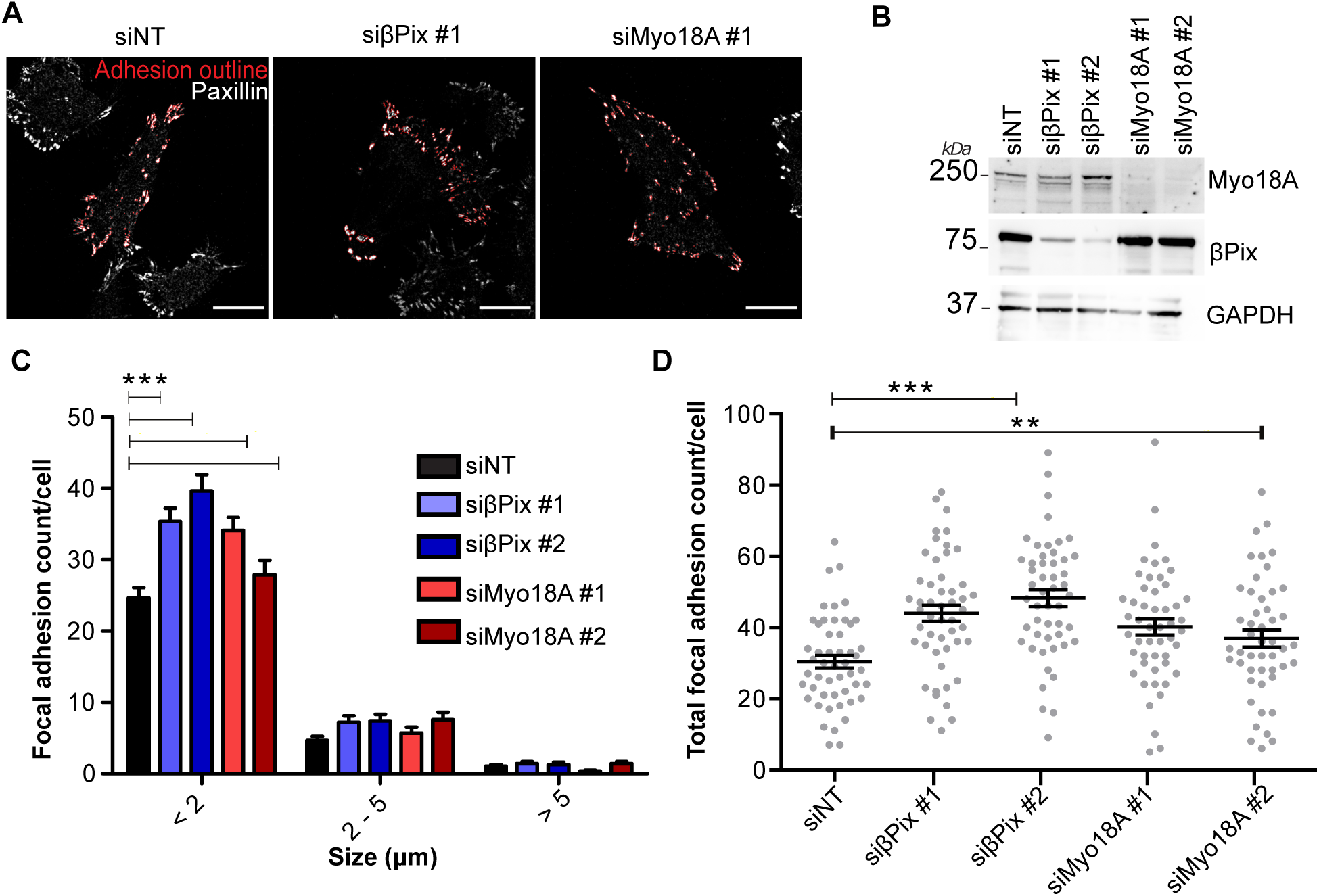
Loss of βPix and Myo18A leads to an increase in nascent adhesions. (A) Representative images of MDA-MB-231 cells non-targeting- (NT), βPix-, or Myo18A-siRNA. fixed and stained with Paxillin. Red mask shows adhesion regions of interest outlined. Scale bar = 10 µm. (B) Representative Western blot of MDA-MB-231 cells treated with non-targeting- (NT), βPix-, or Myo18A-siRNA. Western blot probed for Myo18A, βPix-, and GAPDH (loading control). (C) Histogram showing distribution of adhesion size based on adhesion number. Bonferroni corrected t-tests conducted within each bin comparing each NT-siRNA compared to βPix-, or Myo18A-siRNA treated.(D) Total adhesion count per cell. One-way ANOVA (F(4, 243) = 9.422, P < 0.0001). Dunnett’s Multiple comparison post-test vs. siNT. (N = 5 (pooled), n = 48-50).

**Supplementary Figure 4.**
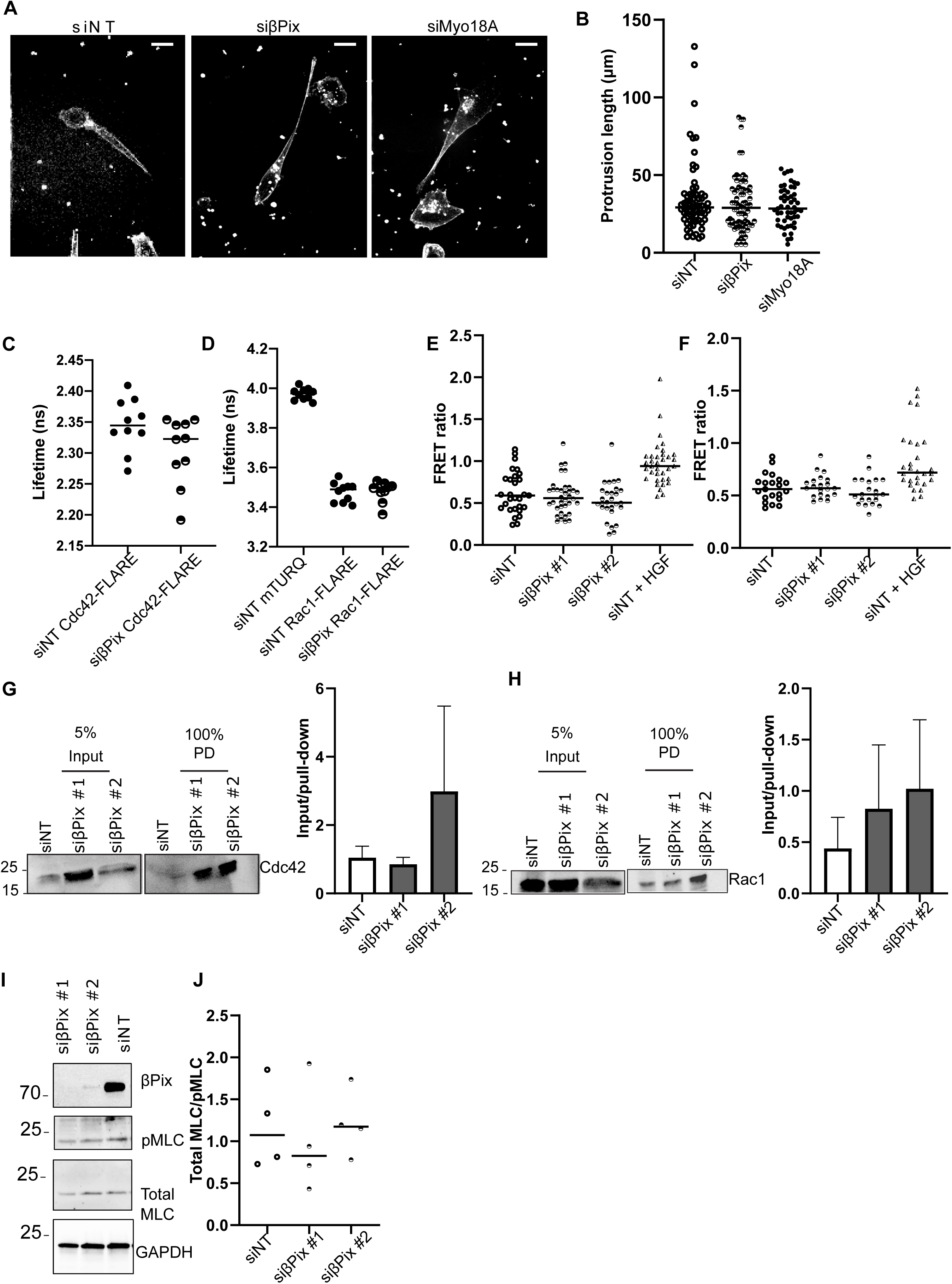
βPix or Myo18A depletion does not affect protrusion assembly, Rho GTPase signalling or MLC phosphorylation. (A) Representative images of MDA-MB-231 cells treated with non-targeting- (NT), βPix-, or Myo18A-siRNA embedded in collagen/fibronectin gels and stained with Sir-Actin. Scale bar = 20 µm. (B) Dot plot shows protrusion length of NT-, βPix-, or Myo18A siRNA-treated cells. Data represents N = 35-40 cells per condition, N = 3 independent experiments. (C) and (D) MDA-MB-231 cells transfected with non-targeting- (NT) or βPix-siRNA, and transfected with Rac- or CDC42-FLARE constructs respectively n = 10 cells, N = 3 independent experiments. (E) and (F) MDA-MB-231 cells transfected with non-targeting- (NT) or βPix-siRNA #1 or βPix-siRNA #2, and transfected with Rac or CDC42 FRET constructs respectively n = 28-35, cells in N = 3. (G) Rac1 and (H) Cdc42 levels in MDA-MB-231 cells transfected with NT- or βPix-siRNA. Cells were lysed 96 h after first siRNA treatment, and Rac1 or Cdc42 were pulled down with GST-PBD-Sepharose beads. Equal amounts of protein were subjected to western blot analysis, which were probed for anti-Rac1 and anti-Cdc42. (I) Western blot analysis of Whole cell lysates from MDA-MB-231 cells treated with NT-siRNA, βPix-siRNA #1 or βPix-siRNA #2 were probed for βPix, pMLC, total MLC, or GAPDH. (J) Quantification of pMLC/MLC levels from cell lysates as in I. Figure corresponds to Figure 2.

**Supplementary Figure 5.**
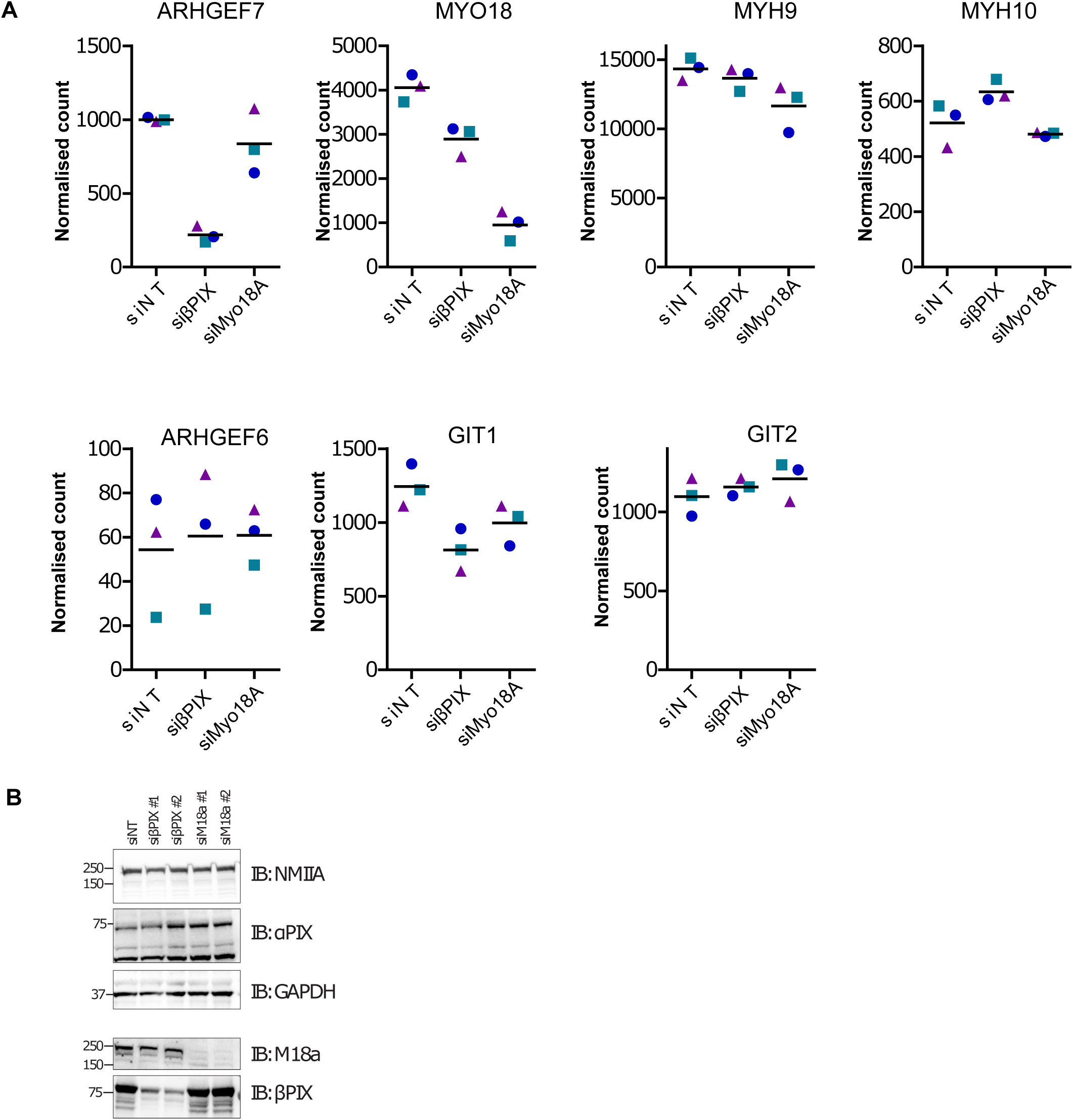
RNA seq analysis and western blot analysis following βPix- or Myo18A-depletion. (A) Normalised counts of selected genes from RNA-Sequencing analysis of MDA-MB-231 cells embedded in collagen/fibronectin gels treated with pooled siRNA oligos: non-targeting- (NT), βPix-, or Myo18A-siRNA (N = 3, Coloured shapes represent individual replicates. ARHGEF7 - βPix protein, MYO18 - Myo18A protein, MYH9 - NM2A heavy chain protein, MYH10 - NM2B heavy chain protein, ARHGEF6 - αPix protein, GIT1 - GIT1 protein, GIT2 - GIT2 protein. (B) Western blot analysis of MDA-MB-231 cells embedded in 3D Collagen treated with pooled siRNA oligos. Figure corresponds to Figure 2.

**Supplementary Figure 6.**
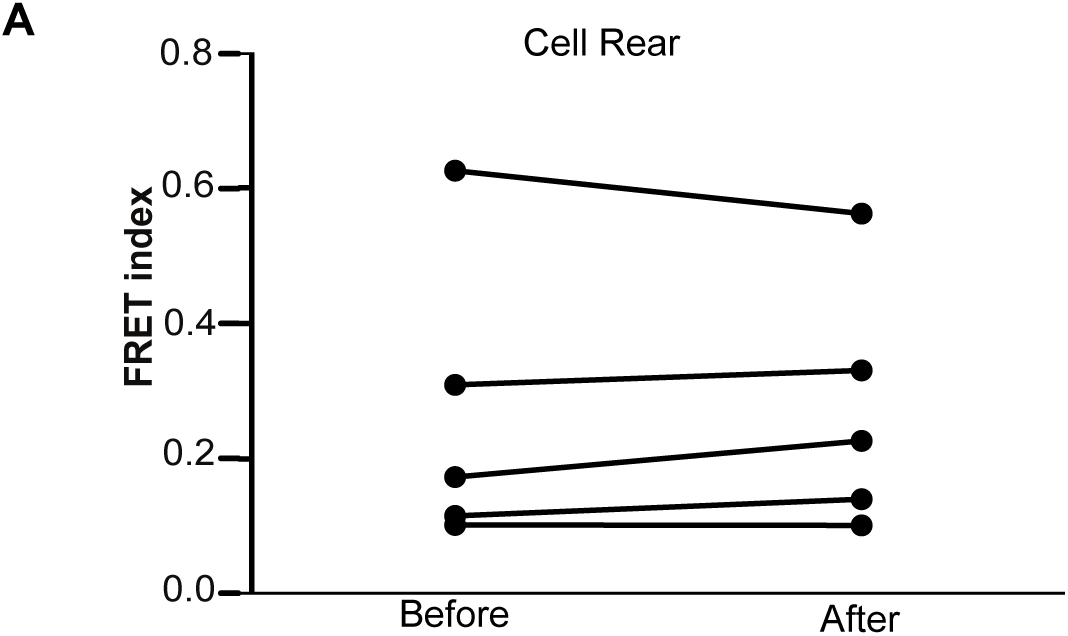
Nuclear tension measured by mini-Nesprin-2 tension sensor. (A) Confocal imaging of MDA-MB-231 cells expressing mN2-TS (mini-Nesprin2 tension sensor) in 3D environments (2mg/mL). Average FRET index of the rear face of the nucleus before and after translocation for 5 cells. P = 0.741, paired t-test. Figure corresponds to Figure 4.

**Supplementary Figure 7.**
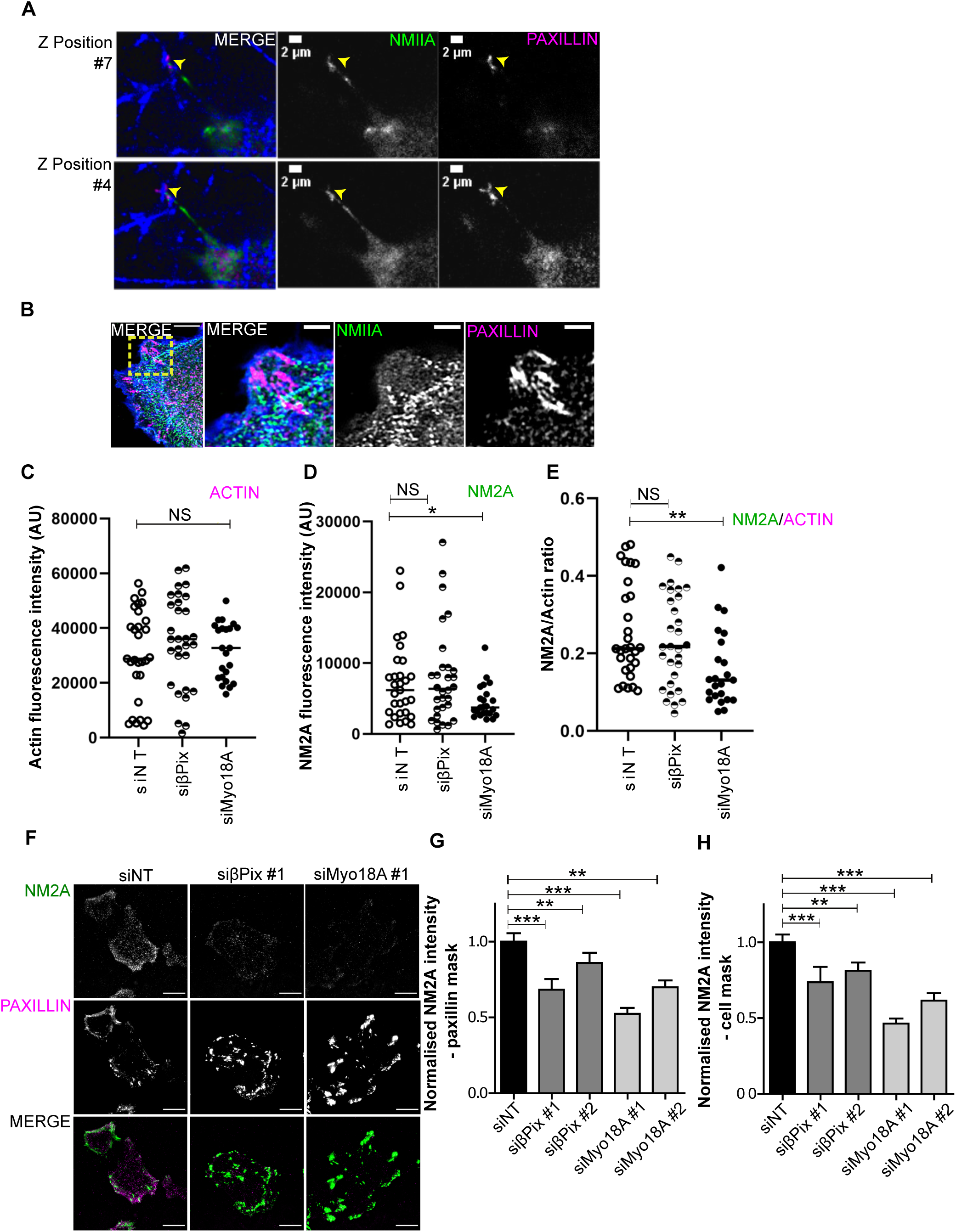
NM2A accumulates at adhesive sites in 3D-embedded cells. (A) CRISPR-generated NM2A-mNeon (green) MDA-MB-231 knock-in cells embedded in collagen (supplemented with ∼1-2% fluorescent collagen, blue) and fibronectin gels were fixed and stained for Paxillin (magenta). Images represent single Z slice projections at two different positions. Adhesive sites are indicated with yellow arrowhead. Corresponds to Figure 6. (B) CRISPR-generated NM2A-mNeon (green) MDA-MB-231 knock-in cells plated on fibronectin/collagen-coated coverslips were fixed and stained for Paxillin (magenta) and phalloidin (blue). Image on left panel is expanded view, yellow boxed region indicate zoom region shown on right. Scale bar expanded = 5 μm, zoom = 2 μm. (C-E) NM2A-mNeon (green) MDA-MB-231 knock-in cells treated with non-targeting- (NT), βPix-, or Myo18A-siRNA embedded in collagen/fibronectin gels were stained with Sir-Actin and imaged by confocal microscopy. Dot-plots show fluorescence intensity of actin (C) and NM2A (D) along the length of the actin protrusion was measured. (E) Ratio of actin:NM2A intensity calculated from data shown in B,C. n = 23-30 cells, N = 3 independent experiments. Mann Whitney test was used to estimate p values: NS - not significant, ** p < 0.01, * p < 0.05. Figure corresponds to Figure 5. (F) NM2A staining on 2D surfaces. Representative TIRF images of MDA-MD-231 cells treated non-targeting- (NT) or βPix-siRNA plated on fibronectin/collagen-coated coverslips fixed and stained for NM2A (green) and Paxillin (magenta). Scale bar = 10 μm. (G) Average NM2A intensity within Paxillin based adhesion mask. One-way ANOVA (F(4,242) = 10.10, P < 0.0001). Dunnett’s multiple comparison post-test vs siNT. (N = 5 (Pooled),n=48−50).

Supplementary Movie 1: **NM2A accumulates early in new protrusions**

MDA-231-2A cell expressing NM2A-mNeon (green) cells transfected with mApple-paxillin (magenta) embedded in collagen supplemented with 1-2% fluorescent collagen (blue)) & fibronectin gels. Cells were imaged by live cell microscopy at 2 min intervals, Scale bar 2 μm. Corresponds to Figure 5.

## References

Aghazadeh, B., Zhu, K., Kubiseski, T. J., Liu, G. A., Pawson, T., Zheng, Y., and Rosen, M. K. (1998). Structure and mutagenesis of the Dbl homology domain. Nat Struct Biol 5, 1098–1107

Ahn, S. J., Chung, K. W., Lee, R. A., Park, I. A., Lee, S. H., Park, D. E., and Noh, D. Y. (2003). Overexpression of betaPix-a in human breast cancer tissues. Cancer Lett 193, 99–107.

Ajeian, J. N., Horton, E. R., Astudillo, P., Byron, A., Askari, J. A., Millon-Frémillon, A., Knight, D., Kimber, S. J., Humphries, M. J., and Humphries, J. D. (2016). Proteomic analysis of integrin-associated complexes from mesenchymal stem cells. Proteomics Clinical applications 10, 51–57.

Alieva, N. O., Efremov, A. K., Hu, S., Oh, D., Chen, Z., Natarajan, M., Ong, H. T., Jégou, A., Romet-Lemonne, G., Groves, J. T., et al. (2019). Myosin IIA and formin dependent mechanosensitivity of filopodia adhesion. Nature Communications 10, 3593.

Arsenovic, Paul T., Ramachandran, I., Bathula, K., Zhu, R., Narang, Jiten D., Noll, Natalie A., Lemmon, Christopher A., Gundersen, Gregg G., and Conway, Daniel E. (2016). Nesprin-2G, a Component of the Nuclear LINC Complex, Is Subject to Myosin-Dependent Tension. Biophysical Journal 110, 34–43.

Barua, B., Nagy, A., Sellers, J. R., and Hitchcock-DeGregori, S. E. (2014). Regulation of nonmuscle myosin II by tropomyosin. Biochemistry 53, 4015–4024.

Billington, N., Beach, Jordan R., Heissler, Sarah M., Remmert, K., Guzik-Lendrum, S., Nagy, A., Takagi, Y., Shao, L., Li, D., Yang, Y., et al. (2015). Myosin 18A Coassembles with Nonmuscle Myosin 2 to Form Mixed Bipolar Filaments. Current Biology 25, 942–948.

Bouzid, T., Kim, E., Riehl, B. D., Esfahani, A. M., Rosenbohm, J., Yang, R., Duan, B., and Lim, J. Y. (2019). The LINC complex, mechanotransduction, and mesenchymal stem cell function and fate. Journal of biological engineering 13, 68–68.

Brown, M. C., and Turner, C. E. (2004). Paxillin: Adapting to Change. Physiological Reviews 84, 1315.

Caswell, P. T., and Zech, T. (2018). Actin-Based Cell Protrusion in a 3D Matrix. Trends Cell Biol 28, 823–834.

Chang, W., Antoku, S., Östlund, C., Worman, H. J., and Gundersen, G. G. (2015). Linker of nucleoskeleton and cytoskeleton (LINC) complex-mediated actin-dependent nuclear positioning orients centrosomes in migrating myoblasts. Nucleus (Austin, Tex) 6, 77–88.

Chastney, M. R., Lawless, C., Humphries, J. D., Warwood, S., Jones, M. C., Knight, D., Jorgensen, C., and Humphries, M. J. (2020). Topological features of integrin adhesion complexes revealed by multiplexed proximity biotinylation. J Cell Biol 219.

Ciobanasu, C., Faivre, B., and Le Clainche, C. (2012). Actin Dynamics Associated with Focal Adhesions. International Journal of Cell Biology 2012, 941292.

Cukierman, E., Pankov, R., Stevens, D. R., and Yamada, K. M. (2001). Taking Cell-Matrix Adhesions to the Third Dimension. Science 294, 1708.

Davidson, P. M., Battistella, A., Déjardin, T., Betz, T., Plastino, J., Borghi, N., Cadot, B., and Sykes, C. (2020). Nesprin-2 accumulates at the front of the nucleus during confined cell migration. EMBO reports 21, e49910.

Davidson, P. M., and Lammerding, J. (2014). Broken nuclei--lamins, nuclear mechanics, and disease. Trends in cell biology 24, 247–256.

DeMali, K. A., Barlow, C. A., and Burridge, K. (2002). Recruitment of the Arp2/3 complex to vinculin: coupling membrane protrusion to matrix adhesion. J Cell Biol 159, 881–891.

Dong, J. M., Tay, F. P., Swa, H. L., Gunaratne, J., Leung, T., Burke, B., and Manser, E. (2016). Proximity biotinylation provides insight into the molecular composition of focal adhesions at the nanometer scale. Sci Signal 9, rs4.

Fraley, S. I., Feng, Y., Krishnamurthy, R., Kim, D.-H., Celedon, A., Longmore, G. D., and Wirtz, D. (2010). A distinctive role for focal adhesion proteins in three-dimensional cell motility. Nature cell biology 12, 598–604.

Goult, B. T., Yan, J., and Schwartz, M. A. (2018). Talin as a mechanosensitive signaling hub. Journal of Cell Biology 217, 3776–3784.

Grashoff, C., Hoffman, B. D., Brenner, M. D., Zhou, R., Parsons, M., Yang, M. T., McLean, M. A., Sligar, S. G., Chen, C. S., Ha, T., and Schwartz, M. A. (2010). Measuring mechanical tension across vinculin reveals regulation of focal adhesion dynamics. Nature 466, 263–266.

Harada, T., Swift, J., Irianto, J., Shin, J. W., Spinler, K. R., Athirasala, A., Diegmiller, R., Dingal, P. C., Ivanovska, I. L., and Discher, D. E. (2014). Nuclear lamin stiffness is a barrier to 3D migration, but softness can limit survival. J Cell Biol 204, 669–682.

Harunaga, J. S., and Yamada, K. M. (2011). Cell-matrix adhesions in 3D. Matrix biology : journal of the International Society for Matrix Biology 30, 363–368.

Hieda, M. (2019). Signal Transduction across the Nuclear Envelope: Role of the LINC Complex in Bidirectional Signaling. Cells 8, 124.

Hiroyasu, S., Stimac, G. P., Hopkinson, S. B., and Jones, J. C. R. (2017). Loss of β-PIX inhibits focal adhesion disassembly and promotes keratinocyte motility via myosin light chain activation. Journal of Cell Science 130, 2329.

Hodgson, L., Shen, F., and Hahn, K. (2010). Biosensors for characterizing the dynamics of rho family GTPases in living cells. Current protocols in cell biology Chapter 14, Unit-14.11.26.

Horton, E. R., Byron, A., Askari, J. A., Ng, D. H. J., Millon-Frémillon, A., Robertson, J., Koper, E. J., Paul, N. R., Warwood, S., Knight, D., et al. (2015). Definition of a consensus integrin adhesome and its dynamics during adhesion complex assembly and disassembly. Nat Cell Biol 17, 1577–1587.

Horton, E. R., Humphries, J. D., James, J., Jones, M. C., Askari, J. A., and Humphries, M. J. (2016). The integrin adhesome network at a glance. Journal of cell science 129, 4159–4163.

Hsu, R.-M., Tsai, M.-H., Hsieh, Y.-J., Lyu, P.-C., and Yu, J.-S. (2010). Identification of MYO18A as a novel interacting partner of the PAK2/betaPIX/GIT1 complex and its potential function in modulating epithelial cell migration. Mol Biol Cell 21, 287–301.

Hsu, R. M., Hsieh, Y. J., Yang, T. H., Chiang, Y. C., Kan, C. Y., Lin, Y. T., Chen, J. T., and Yu, J. S. (2014). Binding of the extreme carboxyl-terminus of PAK-interacting exchange factor β (βPIX) to myosin 18A (MYO18A) is required for epithelial cell migration. Biochim Biophys Acta 1843, 2513–2527.

Hu, S., Dasbiswas, K., Guo, Z., Tee, Y. H., Thiagarajan, V., Hersen, P., Chew, T. L., Safran, S. A., Zaidel-Bar, R., and Bershadsky, A. D. (2017). Long-range self-organization of cytoskeletal myosin II filament stacks. Nat Cell Biol 19, 133–141.

Humphries, J. D., Byron, A., Bass, M. D., Craig, S. E., Pinney, J. W., Knight, D., and Humphries, M. J. (2009). Proteomic analysis of integrin-associated complexes identifies RCC2 as a dual regulator of Rac1 and Arf6. Science signaling 2, ra51-ra51.

Itoh, R. E., Kurokawa, K., Ohba, Y., Yoshizaki, H., Mochizuki, N., and Matsuda, M. (2002). Activation of Rac and Cdc42 Video Imaged by Fluorescent Resonance Energy Transfer-Based Single-Molecule Probes in the Membrane of Living Cells. Molecular and Cellular Biology 22, 6582–6591.

Janota, C. S., Calero-Cuenca, F. J., and Gomes, E. R. (2020). The role of the cell nucleus in mechanotransduction. Current Opinion in Cell Biology 63, 204–211.

Jones, M. C., Humphries, J. D., Byron, A., Millon-Frémillon, A., Robertson, J., Paul, N. R., Ng, D. H. J., Askari, J. A., and Humphries, M. J. (2015). Isolation of Integrin-Based Adhesion Complexes. Current Protocols in Cell Biology 66, 9.8.1–9.8.15.

Kanchanawong, P., Shtengel, G., Pasapera, A. M., Ramko, E. B., Davidson, M. W., Hess, H. F., and Waterman, C. M. (2010). Nanoscale architecture of integrin-based cell adhesions. Nature 468, 580–584.

Kim, D. I., Jensen, S. C., Noble, K. A., Kc, B., Roux, K. H., Motamedchaboki, K., and Roux, K. J. (2016). An improved smaller biotin ligase for BioID proximity labeling. Mol Biol Cell 27, 1188–1196.

Klapholz, B., and Brown, N. H. (2017). Talin - the master of integrin adhesions. J Cell Sci 130, 2435–2446.

Kovács, M., Thirumurugan, K., Knight, P. J., and Sellers, J. R. (2007). Load-dependent mechanism of nonmuscle myosin 2. Proceedings of the National Academy of Sciences 104, 9994–9999.

Kovács, M., Wang, F., Hu, A., Zhang, Y., and Sellers, J. R. (2003). Functional Divergence of Human Cytoplasmic Myosin II: KINETIC CHARACTERIZATION OF THE NON-MUSCLE IIA ISOFORM. Journal of Biological Chemistry 278, 38132–38140.

Kuo, J.-C., Han, X., Hsiao, C.-T., Yates, J. R., 3rd, and Waterman, C. M. (2011). Analysis of the myosin-II-responsive focal adhesion proteome reveals a role for β-Pix in negative regulation of focal adhesion maturation. Nature cell biology 13, 383–393.

Kutys, M. L., and Yamada, K. M. (2014). An extracellular-matrix-specific GEF-GAP interaction regulates Rho GTPase crosstalk for 3D collagen migration. Nature cell biology 16, 909–917.

Lee, J. S. H., Hale, C. M., Panorchan, P., Khatau, S. B., George, J. P., Tseng, Y., Stewart, C. L., Hodzic, D., and Wirtz, D. (2007). Nuclear lamin A/C deficiency induces defects in cell mechanics, polarization, and migration. Biophysical journal 93, 2542–2552.

Livne, A., and Geiger, B. (2016). The inner workings of stress fibers − from contractile machinery to focal adhesions and back. Journal of Cell Science 129, 1293–1304.

Lomakin, A. J., Cattin, C. J., Cuvelier, D., Alraies, Z., Molina, M., Nader, G. P. F., Srivastava, N., Sáez, P. J., Garcia-Arcos, J. M., Zhitnyak, I. Y., et al. (2020). The nucleus acts as a ruler tailoring cell responses to spatial constraints. Science 370, eaba2894.

Lombardi, M. L., Jaalouk, D. E., Shanahan, C. M., Burke, B., Roux, K. J., and Lammerding, J. (2011). The interaction between nesprins and sun proteins at the nuclear envelope is critical for force transmission between the nucleus and cytoskeleton. The Journal of biological chemistry 286, 26743–26753.

Love, M. I., Huber, W., and Anders, S. (2014). Moderated estimation of fold change and dispersion for RNA-seq data with DESeq2. Genome Biology 15, 550.

Luxton, G. W. G., Gomes, E. R., Folker, E. S., Vintinner, E., and Gundersen, G. G. (2010). Linear arrays of nuclear envelope proteins harness retrograde actin flow for nuclear movement. Science (New York, NY) 329, 956–959.

Makowska, Katarzyna A., Hughes, Ruth E., White, Kathryn J., Wells, Claire M., and Peckham, M. (2015). Specific Myosins Control Actin Organization, Cell Morphology, and Migration in Prostate Cancer Cells. Cell Reports 13, 2118–2125.

Manser, E., Loo, T.-H., Koh, C.-G., Zhao, Z.-S., Chen, X.-Q., Tan, L., Tan, I., Leung, T., and Lim, L. (1998). PAK Kinases Are Directly Coupled to the PIX Family of Nucleotide Exchange Factors. Molecular Cell 1, 183–192.

Matellan, C., and del Río Hernández, A. E. (2019). Engineering the cellular mechanical microenvironment – from bulk mechanics to the nanoscale. Journal of Cell Science 132, jcs229013.

Mellacheruvu, D., Wright, Z., Couzens, A. L., Lambert, J. P., St-Denis, N. A., Li, T., Miteva, Y. V., Hauri, S., Sardiu, M. E., Low, T. Y., et al. (2013). The CRAPome: a contaminant repository for affinity purification-mass spectrometry data. Nat Methods 10, 730–736.

Müller, P. M., Rademacher, J., Bagshaw, R. D., Wortmann, C., Barth, C., van Unen, J., Alp, K. M., Giudice, G., Eccles, R. L., Heinrich, L. E., et al. (2020). Systems analysis of RhoGEF and RhoGAP regulatory proteins reveals spatially organized RAC1 signalling from integrin adhesions. Nature Cell Biology 22, 498–511.

Nakade, S., Mochida, K., Kunii, A. et al. Biased genome editing using the local accumulation of DSB repair molecules system. Nat Commun 9, 3270 (2018).

Oh, W. K., Yoo, J. C., Jo, D., Song, Y. H., Kim, M. G., and Park, D. (1997). Cloning of a SH3 Domain-Containing Proline-Rich Protein, p85SPR, and Its Localization in Focal Adhesion. Biochemical and Biophysical Research Communications 235, 794–798.

Owen, L. M., Adhikari, A. S., Patel, M., Grimmer, P., Leijnse, N., Kim, M. C., Notbohm, J., Franck, C., and Dunn, A. R. (2017). A cytoskeletal clutch mediates cellular force transmission in a soft, three-dimensional extracellular matrix. Mol Biol Cell 28, 1959–1974.

Raab, M., and Discher, D. E. (2017). Matrix rigidity regulates microtubule network polarization in migration. Cytoskeleton 74, 114–124.

Rai, V., Thomas, D. G., Beach, J. R., and Egelhoff, T. T. (2017). Myosin IIA Heavy Chain Phosphorylation Mediates Adhesion Maturation and Protrusion in Three Dimensions. The Journal of biological chemistry 292, 3099–3111.

Ridley, A. J., Schwartz, M. A., Burridge, K., Firtel, R. A., Ginsberg, M. H., Borisy, G., Parsons, J. T., and Horwitz, A. R. (2003). Cell Migration: Integrating Signals from Front to Back. Science 302, 1704.

Sakuma, T., Nakade, S., Sakane, Y., Suzuki, K. T., and Yamamoto, T. (2016). MMEJ-assisted gene knock-in using TALENs and CRISPR-Cas9 with the PITCh systems. Nat Protoc 11, 118–133.

Schiller, H. B., Friedel, C. C., Boulegue, C., and Fässler, R. (2011). Quantitative proteomics of the integrin adhesome show a myosin II-dependent recruitment of LIM domain proteins. EMBO reports 12, 259–266.

Seetharaman, S., and Etienne-Manneville, S. (2020). Cytoskeletal Crosstalk in Cell Migration. Trends in Cell Biology 30, 720–735.

Senger, F., Pitaval, A., Ennomani, H., Kurzawa, L., Blanchoin, L., and Théry, M. (2019). Spatial integration of mechanical forces by α-actinin establishes actin network symmetry. Journal of Cell Science 132, jcs236604.

Serrels, B., Serrels, A., Brunton, V. G., Holt, M., McLean, G. W., Gray, C. H., Jones, G. E., and Frame, M. C. (2007). Focal adhesion kinase controls actin assembly via a FERM-mediated interaction with the Arp2/3 complex. Nat Cell Biol 9, 1046–1056.

Shutova, M. S., Asokan, S. B., Talwar, S., Assoian, R. K., Bear, J. E., and Svitkina, T. M. (2017). Self-sorting of nonmuscle myosins IIA and IIB polarizes the cytoskeleton and modulates cell motility. Journal of Cell Biology 216, 2877–2889.

Soneson, C., Love, M., and Robinson, M. (2015). Differential analyses for RNA-seq: transcript-level estimates improve gene-level inferences [version 1; peer review: 2 approved]. F1000Research 4.

Thomas, D. G., Yenepalli, A., Denais, C. M., Rape, A., Beach, J. R., Wang, Y.-L., Schiemann, W. P., Baskaran, H., Lammerding, J., and Egelhoff, T. T. (2015). Non-muscle myosin IIB is critical for nuclear translocation during 3D invasion. The Journal of cell biology 210, 583–594.

Tojkander, S., Gateva, G., and Lappalainen, P. (2012). Actin stress fibers – assembly, dynamics and biological roles. Journal of Cell Science 125, 1855–1864.

van Helvert, S., Storm, C., and Friedl, P. (2018). Mechanoreciprocity in cell migration. Nature Cell Biology 20, 8–20.

Vicente-Manzanares, M., Newell-Litwa, K., Bachir, A. I., Whitmore, L. A., and Horwitz, A. R. (2011). Myosin IIA/IIB restrict adhesive and protrusive signaling to generate front-back polarity in migrating cells. The Journal of cell biology 193, 381–396.

Vignaud, T., Copos, C., Leterrier, C., Toro-Nahuelpan, M., Tseng, Q., Mahamid, J., Blanchoin, L., Mogilner, A., Théry, M., and Kurzawa, L. (2021). Stress fibres are embedded in a contractile cortical network. Nat Mater 20, 410–420.

Wang, F., Kovács, M., Hu, A., Limouze, J., Harvey, E. V., and Sellers, J. R. (2003). Kinetic Mechanism of Non-muscle Myosin IIB: FUNCTIONAL ADAPTATIONS FOR TENSION GENERATION AND MAINTENANCE. Journal of Biological Chemistry 278, 27439–27448.

Wolf, K., Te Lindert, M., Krause, M., Alexander, S., Te Riet, J., Willis, A. L., Hoffman, R. M., Figdor, C. G., Weiss, S. J., and Friedl, P. (2013). Physical limits of cell migration: control by ECM space and nuclear deformation and tuning by proteolysis and traction force. The Journal of cell biology 201, 1069–1084.

Woroniuk, A., Porter, A., White, G., Newman, D. T., Diamantopoulou, Z., Waring, T., Rooney, C., Strathdee, D., Marston, D. J., Hahn, K. M., et al. (2018). STEF/TIAM2-mediated Rac1 activity at the nuclear envelope regulates the perinuclear actin cap. Nature Communications 9, 2124.

Young, L. E., and Higgs, H. N. (2018). Focal Adhesions Undergo Longitudinal Splitting into Fixed-Width Units. Current biology : CB 28, 2033–2045.e2035.

Yu, X., Zech, T., McDonald, L., Gonzalez, E. G., Li, A., Macpherson, I., Schwarz, J. P., Spence, H., Futó, K., Timpson, P., et al. (2012). N-WASP coordinates the delivery and F-actin-mediated capture of MT1-MMP at invasive pseudopods. The Journal of cell biology 199, 527–544.

